# Tamm-Horsfall protein augments neutrophil NETosis during urinary tract infection

**DOI:** 10.1101/2024.02.01.578501

**Authors:** Vicki Mercado-Evans, Claude Chew, Camille Serchejian, Alexander Saltzman, Marlyd E. Mejia, Jacob J. Zulk, Ingrid Cornax, Victor Nizet, Kathryn A. Patras

## Abstract

Urinary neutrophils are a hallmark of urinary tract infection (UTI), yet the mechanisms governing their activation, function, and efficacy in controlling infection remain incompletely understood. Tamm-Horsfall glycoprotein (THP), the most abundant protein in urine, uses terminal sialic acids to bind an inhibitory receptor and dampen neutrophil inflammatory responses. We hypothesized that neutrophil modulation is an integral part of THP-mediated host protection. In a UTI model, THP-deficient mice showed elevated urinary tract bacterial burdens, increased neutrophil recruitment, and more severe tissue histopathological changes compared to WT mice. Furthermore, THP-deficient mice displayed impaired urinary NETosis during UTI. To investigate the impact of THP on NETosis, we coupled *in vitro* fluorescence-based NET assays, proteomic analyses, and standard and imaging flow cytometry with peripheral human neutrophils. We found that THP increases proteins involved in respiratory chain, neutrophil granules, and chromatin remodeling pathways, enhances NETosis in an ROS-dependent manner, and drives NET-associated morphologic features including nuclear decondensation. These effects were observed only in the presence of a NETosis stimulus and could not be solely replicated with equivalent levels of sialic acid alone. We conclude that THP is a critical regulator of NETosis in the urinary tract, playing a key role in host defense against UTI.

## INTRODUCTION

Urinary tract infections (UTI) impact around 400 million people globally each year, with approximately half of all women experiencing at least one UTI during their lifetime(1–3). The most common culprit of UTIs, responsible for upwards of 75% of cases, is uropathogenic *Escherichia coli* (UPEC)(1, 4, 5). Genetic factors that increase UTI susceptibility include variants in bacterial ligand recognition, innate immune signaling and neutrophil recruitment(6–8). A hallmark clinical feature of UTI is the rapid recruitment of neutrophils following *E. coli* introduction(9, 10) corresponding with the onset of UTI symptoms(11). Murine models demonstrate that successful resolution of UTI requires a robust neutrophil response. Neutrophils, the initial responders to UTI, are detected in urine as early as 2 hours post-infection(12–14). In line with clinical observations of genetic risk factors, aberrant neutrophil recruitment in mice leads to pathological neutrophil accumulation, tissue damage, and scarring(15, 16), whereas antibody-mediated neutrophil depletion exacerbates bacterial burdens and promotes chronic infection(12, 17).

Neutrophils display diverse antibacterial functions that contribute to the resolution of UTI. They are a critical source of antimicrobial proteins including cathelicidin(18, 19) and lactoferrin(20, 21) and are the principal cell type performing bacterial phagocytosis *in vivo*(22). Additionally, neutrophils are a key source of reactive oxygen species, essential for bacterial killing, but which in excess can promote tissue damage, particularly in the kidneys(23, 24). Neutrophils isolated from patients with recurrent UTI display decreased phagocytosis and reduced production of reactive oxygen intermediates underscoring the importance of these functions for neutrophil antibacterial activity and the resolution of UTI(25).

Another neutrophil antimicrobial mechanism, first described in 2004, is NETosis – the process of forming neutrophil extracellular traps (NETs)(26). NETosis is a form of cell death resulting in expulsion of a scaffold of decondensed chromatin studded with antimicrobial products such as myeloperoxidase (MPO), cathelicidin, and histones, that trap various extracellular pathogens to aid in infection control (26, 27). Multiple stimuli can trigger NETosis including phorbol-myristate acetate (PMA, a protein kinase C activator), lipopolysaccharide (LPS), calcium ionophores, hydrogen peroxide, and various microbes, including Gram-positive and Gram-negative bacteria, as well as fungal species(28). Distinct subtypes of NETosis, discriminated based on cellular morphology and signaling pathways, include classical (or suicidal) NETosis(29, 30), mitochondrial NETosis, where NETs are formed from mitochondrial DNA(31), and nonclassical (or vital) NETosis, where the neutrophil expels nuclear DNA without or prior to lysing(32, 33). While recent studies have reported the presence of NET-associated products (e.g. DNA, histones, MPO) in the urine of patients with UTIs(34, 35) and have demonstrated NET formation in UTI using a bladder-on-a-chip model(36), the role of NETosis in UTI susceptibility and clearance remains to be established.

We hypothesized that urinary specific factors may influence the formation of NETs in UTI. Tamm-Horsfall protein (THP), the most abundant urinary protein, is a highly conserved glycoprotein with multiple important roles in urinary tract health including the regulation of salt and water homeostasis and the prevention of mineral crystallization(37, 38). In the context of UTI, THP is a key host defense factor. Elimination of THP increases UTI susceptibility in murine models(39–42). THP directly binds urinary pathogens(42–46), inhibiting microbial adherence to host urothelium, which in turn aids clearance via urinary excretion. THP also shapes host responses to UPEC by modulating immune cell activity in a cell type and context-dependent manner(47–49). We previously showed that THP terminal sialic acids engage Siglec-9, an inhibitory neutrophil receptor, to suppress neutrophil activities including chemotaxis, ROS release, and bactericidal capacity(50). This immunosuppressive impact of THP is revealed in THP KO mice, which exhibit elevated circulating pro-inflammatory cytokines, increased kidney inflammation during renal injury, and neutrophilia in the blood and urine with or without inflammatory stimuli(50–53). However, the modulation of host immune responses by THP in the context of UTI has not been reported.

Given the protective roles of both THP and neutrophils in UTI, and considering THP influence on neutrophil responses, we hypothesized that THP may provide additional host protection by modulating NETosis. To investigate this hypothesis, we evaluated neutrophil recruitment and NETosis in a murine UTI model comparing wildtype to THP-deficient mice. Our findings revealed increased bladder neutrophil recruitment in THP-deficient mice, but reduced NET formation compared to wildtype mice. Subsequent validation through flow cytometry of human neutrophils confirmed that THP enhancement of NETosis was dependent on neutrophil activation and reactive oxygen species. In conjunction with its roles in impeding pathogen adherence and tempering excessive inflammation, we conclude that THP provides added host protection by modulating NETosis during UTI.

## RESULTS

### THP deficiency increases urinary tract UPEC burdens and tissue histopathology

Prior studies have identified the heightened susceptibility of THP-deficient mice to elevated UPEC burdens in the urine and bladder at 24 hours post-infection(39, 40). To assess the sustained impact of THP deficiency during acute UTI, we used an established model of UPEC UTI(54) in THP^+/+^ (WT) and THP^-/-^ (KO) mice(50). In this model, mice receive 1×10^8^ CFU of UPEC cystitis strain UTI89 or are mock-infected as a control. Consistent with previous findings(39, 40), THP KO mice exhibited persistent increased bacteriuria (**Fig. 1A**), and temporarily elevated bacterial load in the bladder and kidney throughout the infection course (**Fig. 1B-C**) compared to WT mice. Bladder and kidney tissue sections collected during acute infection were examined by a blinded veterinary pathologist and scored on a 0-4 scale, considering pathologic features such as intraluminal bacteria, submucosal edema, and suppurative pyelonephritis. UPEC-infected THP KO mice displayed more severe bladder and kidney pathology compared to their WT counterparts (**Fig. 1D-E**), marked by increased immune infiltration of the urinary epithelium and submucosa (**Fig. 1F-G**) and luminal mixed immune cell aggregation in the renal pelvis (**Fig. 1G**). No differences in histopathology scores were observed in mock-infected WT and THP KO mice.

**Figure 1.**
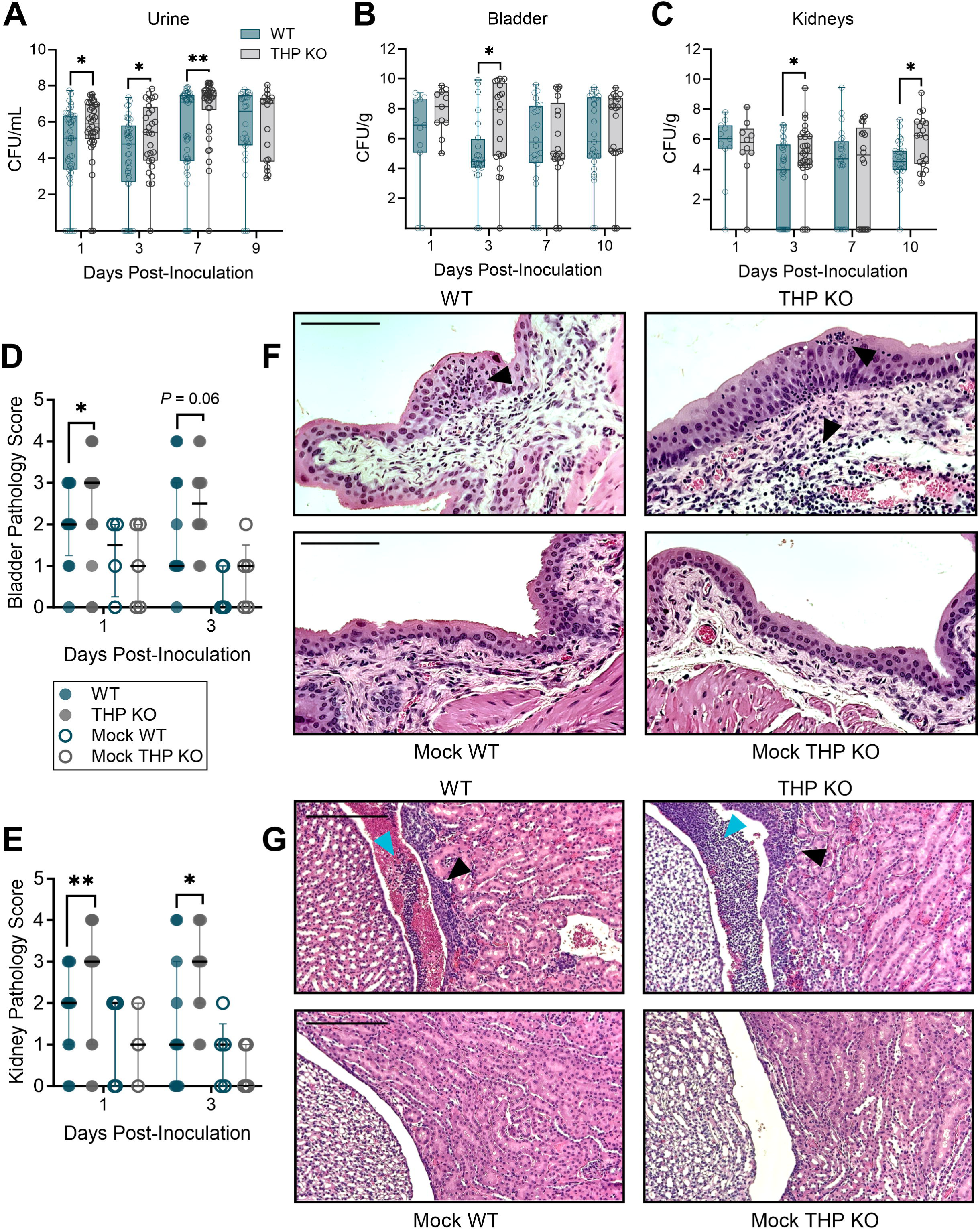
THP deficiency increases urinary tract bacterial burdens and tissue pathology. Wild type (WT) and THP knockout (THP KO) mice were transurethrally infected with 10^8^ CFU of UPEC strain UTI89 or mock-infected as a control. Time course of urine (**A**), bladder(**B**), and kidney (**C**) UPEC burdens from WT and THP KO mice. Bladder (**D**) and kidney (**E**) pathology scores on days 1 and 3 post-infection. (**F**) Representative H&E images of day 1 bladders from UPEC-infected or mock-infected WT and THP KO mice. (**G**) Representative H&E images of day 3 kidneys from UPEC-infected or mock-infected WT and THP KO mice. Scale bars represent 110 µm (F) and 210 µm (G). Black arrows point to polymorphonuclear cell infiltration and blue arrows point to polymorphonuclear cell aggregates in the renal pelvis. Experiments were performed at least two times with data combined. *n* = 18-46/timepoint (A), *n* = 11-31 (B-C), or *n* = 4-15 (D-E). Box and whisker plots extend from 25th to 75th percentiles and show all points (A-C). Points represent individual samples and lines indicate medians (D, E). Data was analyzed by two-tailed Mann-Whitney t-test (A-C) and two-sided Fisher’s exact test (D, E). * *P* < 0.05; ** *P* < 0.01.

### THP deficiency alters bladder neutrophil infiltration and impact of neutrophil depletion during UTI

We next evaluated immune cell infiltration in the bladder and kidneys during UTI by flow cytometry. We surveyed the total immune cell fraction (CD45^+^, P1), as well as neutrophils (Ly6G^+^), non-myeloid (CD11b^−^ CD11c^−^), myeloid (CD11b^±^ CD11c^±^, P3), myeloid antigen presenting cells (APCs, MHC-II), and myeloid non-APC population subsets (gating scheme provided in **Fig. 2A**). At 3 days post-infection, THP KO mice had significantly higher proportions of CD45^+^ cells in both the bladder and kidneys compared to WT-infected mice, although no differences were observed in later timepoints or mock controls (**Fig. 2B-C**). Additionally, bladder neutrophil proportions were elevated infected THP KO mice compared to WT mice at both three and seven days after UPEC inoculation, with no observed differences in the kidneys (**Fig. 2D-E**). Other bladder immune cell sub-populations did not differ between groups (**Supp.** Fig. 1). In the kidneys, minimal differences in other sub-populations were noted including a reduced proportion of myeloid lineages at day 3 and myeloid APCs at day 7 post-inoculation in THP KO compared to WT mice (**Supp.** Fig. 2). Under mock-infected conditions, THP KO mice exhibited a slight but significant increase in the proportion of non-APC myeloid cells (**Supp.** Fig. 2E).

**Figure 2.**
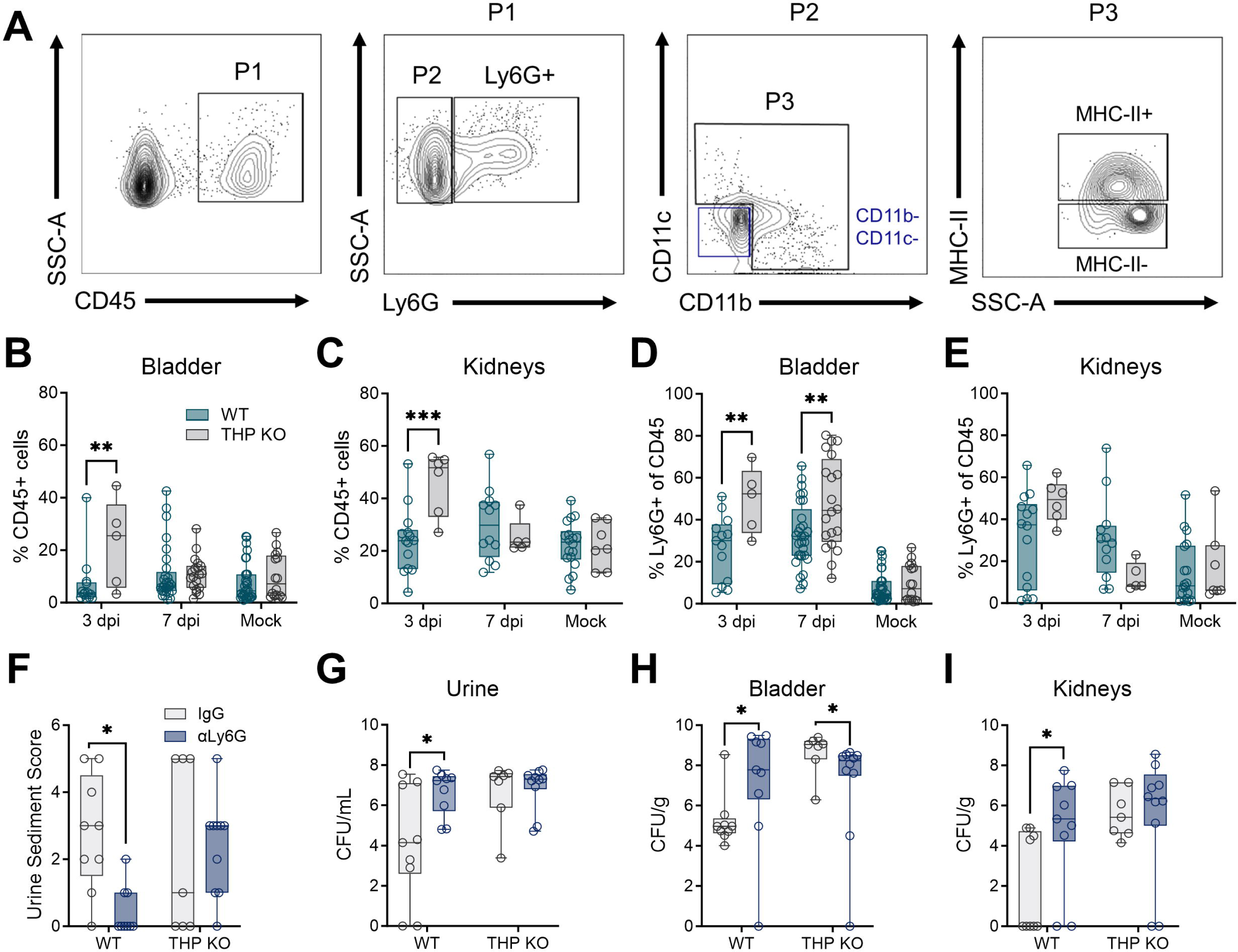
THP deficiency increases bladder neutrophil infiltration and reduces bacterial burdens upon neutrophil depletion during UTI. Wild type (WT) and THP knockout (THP KO) mice were transurethrally infected with 10^8^ CFU of UPEC strain UTI89 or mock-infected as a control. (**A**) Gating strategy for quantifying immune populations of interest with a focus on neutrophils (Ly6G^+^) and myeloid lineages (CD11b±, CD11c±). Frequency of CD45+ cells (P1) cells in bladder (**B**) or kidneys (**C**) in UPEC-challenged or mock-infected THP WT and KO mice 3 and 7 days post-infection (dpi). Mock-infected samples from both timepoints were combined prior to analyses. Frequency of neutrophils (Ly6G+) from CD45+ populations infiltrating bladder (**D**) or kidneys (**E**) in UPEC-challenged or mock-infected mice 3 and 7 days post-infection. To partially deplete neutrophils, mice were administered anti-Ly6G or IgG isotype control just prior to bacterial inoculation and on day 2 and 4 post-inoculation. (**F**) Urine sediment scores at 6 days post-infection. (**G**) Urine UPEC burdens at 6 days post-infection. Bladder (**H**) and kidney (**I**) UPEC burdens at 7 days post-infection. Experiments were performed at least two times with data combined. *n* = 5-32/group (B-E) and *n* = 7-10 (F-I). Box and whisker plots extend from 25th to 75th percentiles and show all points (B-I). Data was analyzed by two-way ANOVA with Sidak’s multiple comparisons test (B-E, G-I) and two-sided Fisher’s exact test (F). * *P* < 0.05; ** *P* < 0.01; *** *P* < 0.001.

Neutrophil depletion has been previously shown to exacerbate bacterial burdens and promote chronic infection depending on the extent of neutrophil reduction(17). To evaluate the impact of neutrophil depletion in THP-deficient mice, mice were administered anti-Ly6G antibody or isotype IgG control intraperitoneally (i.p.) at a dose of 10 μg every 48 h, from day 0 thru day 6 post-inoculation. On day 6, urine sediment was scored for the presence of polymorphonuclear (PMN) cells on a scale of 0 (<1 PMN per high-powered field) to 5 (>40 PMN) as described previously(17). Urine and tissues were collected on day 7 post-inoculation to quantify bacterial burdens. Anti-Ly6G antibody treatment significantly reduced urine sediment PMN scores in WT mice but had no such effect in THP KO mice (**Fig. 2F**). Additionally, anti-Ly6G antibody treatment resulted in increased urine and tissue bacterial burdens in WT mice (**Fig. 2G-I**). In contrast, anti-Ly6G antibody treatment led to decreased bladder CFU in THP KO mice, with no differences observed in urine or kidney burdens.

THP KO mice display heightened inflammation in response to acute kidney injury(52) and show elevated immune cell recruitment during acute UTI (**Fig. 2B-C**). Given that deficiency in cyclooxygenase 2 (COX-2), a critical enzyme initiating inflammatory cascades, downregulates THP expression in the kidneys, and COX-2^-/-^ mice are hyper-susceptible to UTI(55), we examined the impact of COX-2 in our model. Mice were treated with diclofenac, a COX-2 inhibitor(56), provided at 0.2mg/mL in drinking water from day 0 thru day 6 post-inoculation, with tissues collected at day 7. We found no differences in bladder or kidney burdens between diclofenac-treated and mock-treated WT and THP KO mice (**Supp. Fig. S3**). Together, these findings highlight that elevated neutrophils are a distinctive feature of THP deficiency in UTI which, paired with enhanced bacterial burdens, suggest impaired neutrophil activity in THP KO mice.

### Murine urinary THP levels and glycosylation patterns change minimally during UTI

While genetic and clinical studies have linked *UMOD* variants(57) or reduced THP production(58, 59) with enhanced risk for UTI; no differences in urinary THP levels were observed between a pediatric UTI cohort and healthy controls(60). Similarly, we observed no variations in urinary THP levels between mock-infected and UPEC-infected WT mice at days 2-4 post-inoculation (**Fig. 3A**). To delineate the N-glycan profile of murine THP and assess the impact of UTI on THP glycosylation, we collected urine over the first 72 h post-inoculation or mock-treatment and profiled THP glycosylation patterns using MALDI-TOF/TOF mass spectrometry of procainamide-labeled permethylated N-glycans. Similar to human THP(61–63), murine THP contained multiple bi-, tri-, and tetra-antennary sialylated and/or fucosylated complex type N-glycans (**Fig. 3B**). The highest intensity peak (m/z 4588) represented a tetra-antennary, tetra-sialylated and fucosylated N-glycan (**Fig. 3B, Supp. Table 1**) which matches the most abundant N-glycan reported on human THP(61, 62). Other high intensity peaks were observed at m/z 2967, 3777, and 4226. In UPEC-infected THP samples, the four most abundant structures (m/z 2967, 3777, 4226, and 4588) remained the same, albeit with some proportional differences: the m/z 2967 peak, corresponding to a bi-antennary, bi-sialylated and fucosylated N-glycan, increased and the m/z 4588 peak decreased relative to mock-treated spectra (**Fig. 3C, Supp. Table 1**). We quantified the total sialic acid released from purified murine THP by DMB-HPLC analysis. N-glycolylneuraminic acid (Neu5Gc) and N-Acetylneuraminic acid (Neu5Ac) were distinguished by retention times of 4.2 and 5.3 minutes respectively. No differences in Neu5Gc, Neu5Ac, or total sialic acid levels were observed between mock-infected and UPEC-infected samples (**Table 1**). As a control, samples from THP KO mice showed significantly reduced Neu5Ac and total sialic acid compared to WT mock samples validating that THP was the primary source of sialic acid in these analyses (**Table 1**). Together, these data reveal conserved glycosylation patterns, including sialylation by Neu5Ac, in murine and human THP, which are retained during UTI in a mouse model.

**Figure 3.**
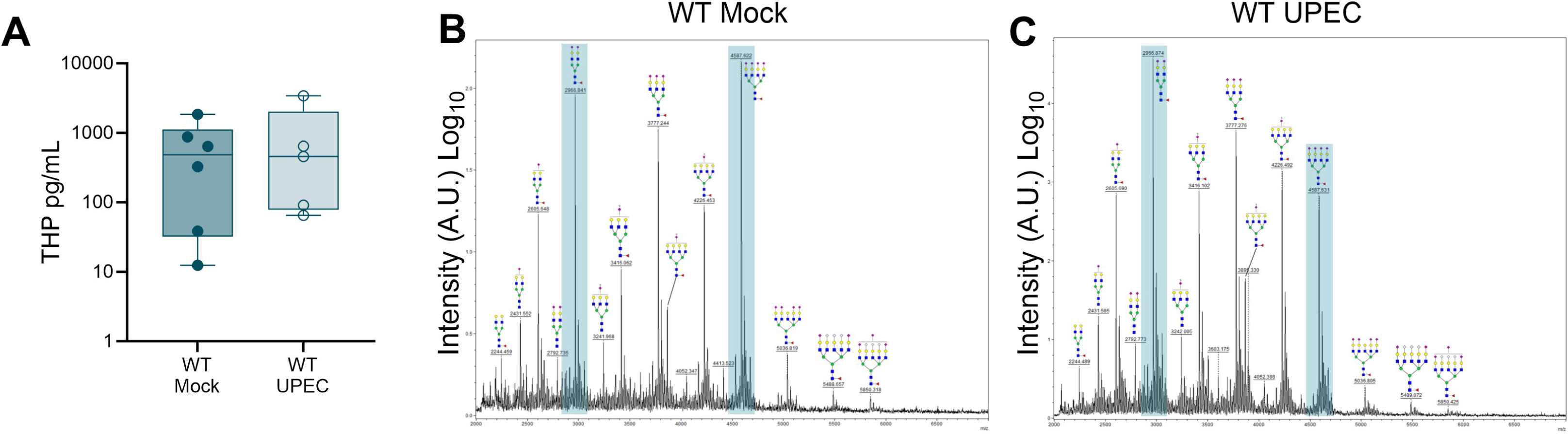
Tamm-Horsfall protein levels and glycosylation patterns change minimally during urinary tract infection *in vivo*. Mouse urine was collected multiple times over the first 4 days post-inoculation from UPEC-infected and mock-infected WT mice. (**A**) THP levels in urine as measured by ELISA. N-glycan MALDI-tof profiles of urinary THP isolated from WT mice that were either mock (**B**) or UPEC-infected (**C**). Data represent one MALDI-tof analysis of purified THP harvested from WT mice (*n* = 15 mock, *n* = 28 UPEC) from two independent experiments. Prominent peaks with proportional differences between UPEC-infected and mock samples (m/z 2967 and m/z 4588) are highlighted in teal. Data (A) was analyzed by two-tailed Mann-Whitney t-test and comparisons were not significant.

**Table 1.**
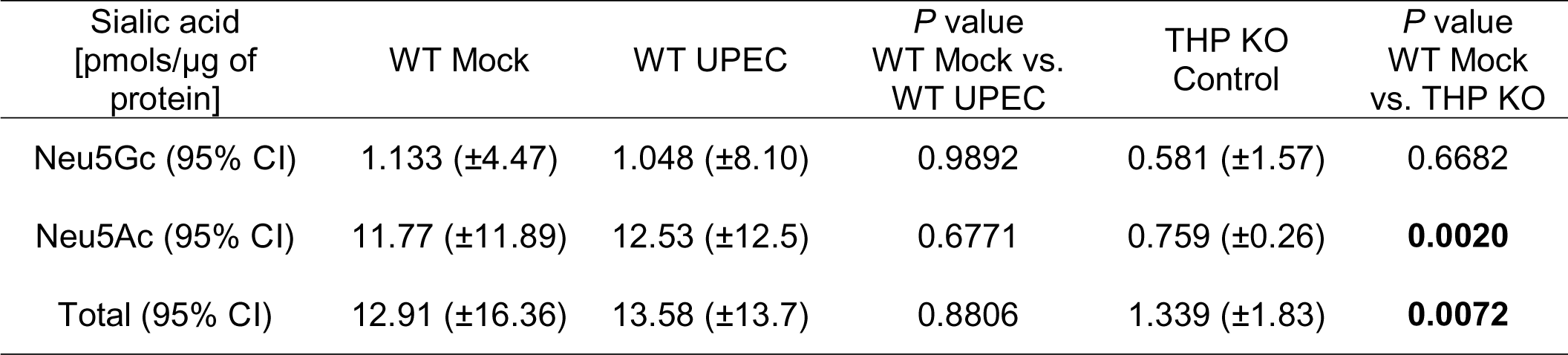
Sialic acid concentration in THP purified from mouse urine. Sialic acid content of THP isolated from pooled mouse urine from WT and THP KO mice as measured by UPLC. Data represent two independent replicate experiments. Data were analyzed by one-way ANOVA with Holmes-Sidak multiple comparison tests.

| Sialic acid<br>[pmols/μg of<br>protein] | WT Mock | WT UPEC | *P* value<br>WT Mock vs.<br>WT UPEC | THP KO<br>Control | *P* value<br>WT Mock<br>vs. THP KO |
| --- | --- | --- | --- | --- | --- |
| Neu5Gc (95% CI) | 1.133 (±4.47) | 1.048 (±8.10) | 0.9892 | 0.581 (±1.57) | 0.6682 |
| Neu5Ac (95% CI) | 11.77 (±11.89) | 12.53 (±12.5) | 0.6771 | 0.759 (±0.26) | **0.0020** |
| Total (95% CI) | 12.91 (±16.36) | 13.58 (±13.7) | 0.8806 | 1.339 (±1.83) | **0.0072** |

### Murine neutrophils undergo NETosis during UTI and THP deficiency alters neutrophil sub-populations

To investigate whether differences in neutrophil abundance (**Fig. 2**) corresponded with differences in neutrophil function, we quantified and visualized NETosis in mouse urine. Nucleic acid dyes including cell-permeable Hoechst 33342 or DAPI and non-cell permeable Sytox dyes have been used to distinguish NETosis from other forms of cell death, including apoptosis, in both human and murine neutrophils in mixed-cell populations(64–66). Additionally, plasma membrane permeability can be confirmed using a live/dead amine-reactive dye that can only fluorescently label intracellular amines if the plasma membrane has been compromised(67). In classical (or suicidal) NETosis, neutrophils permeabilize and expel decondensed chromatin whereas during nonclassical (or vital) NETosis, neutrophils also release DNA but still retain viability and effector functions(68, 69). We collected mouse urine from UPEC-infected or mock-treated mice 24 h post-inoculation, and cells were stained and subjected to flow cytometry. Neutrophils (PMNs) were identified as CD11b^+^ Ly6G^+^ and were further gated based on staining for presence of extracellular DNA (Sytox Orange, SO) as an indicator of NETosis, and plasma membrane permeability (Live/Dead stain) as depicted in **Fig. 4A**. We identified four unique populations: live PMNs (SO^−^ Live/Dead^−^, Q4), dead PMNs (SO^−^ Live/Dead^+^, Q3), classical NETosis (SO^+^ Live/Dead^+^), and nonclassical NETosis (SO^+^ Live/Dead^−^, Q1). WT mice displayed an increase in total urinary neutrophils during infection compared to mock-infected counter parts (**Fig. 4B**). In both WT and THP KO mice, frequency of live PMNs was reduced during infection (**Fig. 4C**), but no significant differences were observed in dead PMNs (**Fig. 4D**). UPEC infection elevated total NETosis (Q1 + Q2) and classical NETosis (Q2) frequency in both WT and THP KO mice compared to their mock-infected counterparts (**Fig. 4E-F**). Uniquely, WT mice showed elevated frequency (**Fig. 4C**) and counts (**Fig. 4D**) of nonclassical NETosis in response to UPEC infection, and compared to UPEC-infected THP KO mice, suggesting that THP promotes nonclassical NETosis during UTI. The presence of NETs in WT and THP KO urine samples at 24 h post-infection was visualized by immunofluorescence microscopy using antibodies for neutrophils (myeloperoxidase, MPO), NETosis (citrullinated histone H3, H3Cit), and THP (**Fig. 4I-J**).

**Figure 4.**
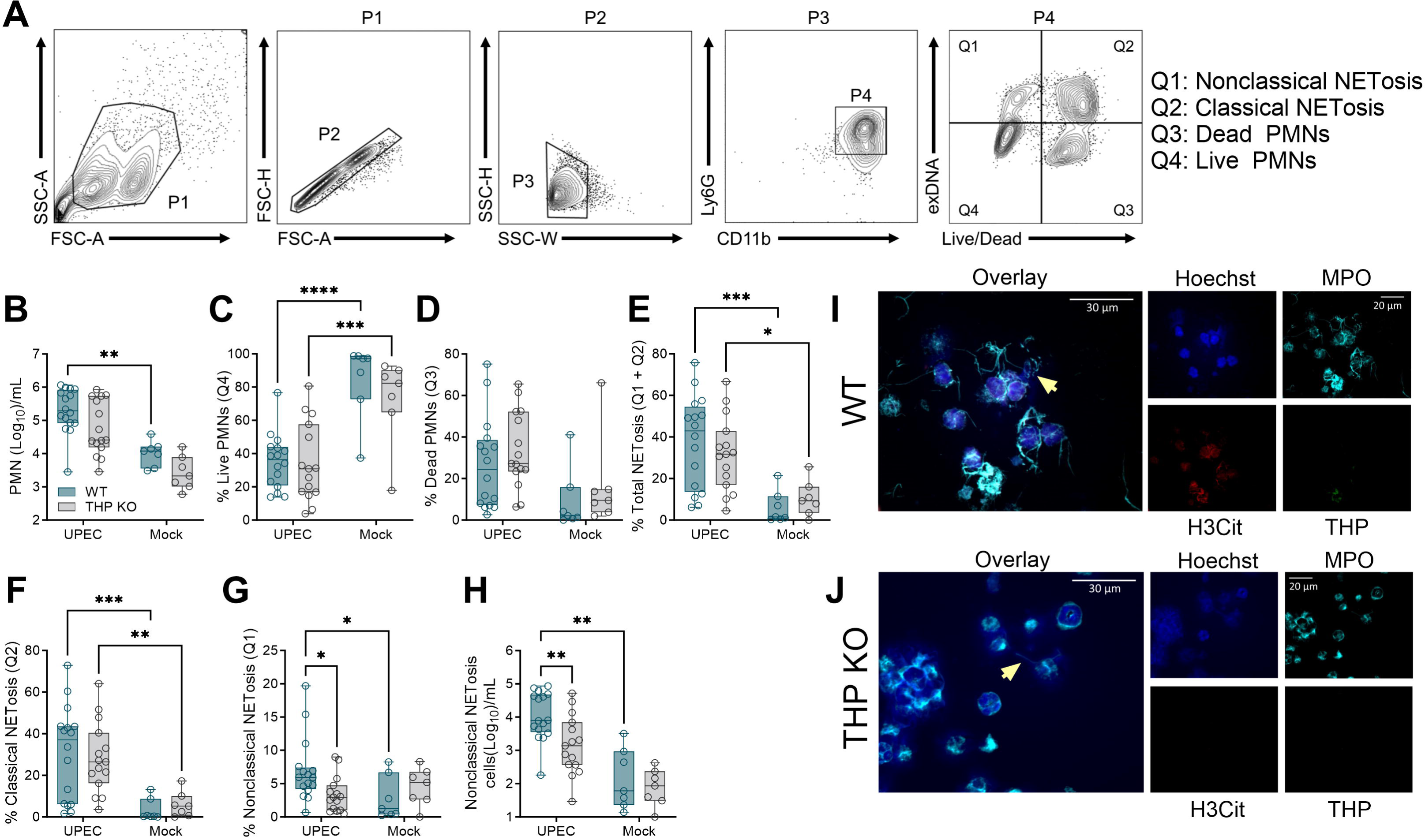
Neutrophil nonclassical NETosis populations are decreased in THP-deficient mice during UTI. Urine was collected from WT and THP KO mice 24 hours post-infection with UPEC or mock-infected as controls. (**A**) Gating strategy for quantifying neutrophil (polymorphonuclear cells, PMNs, Ly6G+, CD11b+, P4) subpopulations of interest with a focus on nonclassical NETosis (extracellular DNA[Sytox Orange]+, Live/Dead-), classical NETosis (exDNA+, Live/Dead+), dead PMNs (exDNA-, Live/Dead+), and live PMNs (exDNA-, Live/Dead-). (**B**) Total PMNs (P4) per mL of urine. Frequency of live PMNs (**C**) and dead PMNs (**D**) out of total PMNs. (**E**) Frequency of total NETosis (Q1 + Q2) out of total PMNs. Frequency of classical NETosis (Q2) (**F**) and nonclassical NETosis (Q1) (**G**) out of total PMNs. (**H**) Nonclassical NETosis cell counts per mL of urine. Urine samples from UPEC-infected WT (**I**) and THP KO (**J**) mice were mounted on slides and NETs were visualized via immunofluorescence using antibodies against myeloperoxidase (MPO, cyan channel), citrullinated histone H3 (H3Cit, red channel), and THP (green channel). Nucleic acids were stained using Hoechst dye (blue channel). Yellow arrows point to NETs structures depicted as strands of DNA dotted with MPO staining. Representative images are shown. Scale bars represent 20 µm (single channels) and 30 µm (inset overlays). Experiments were performed at least two times with data combined. *n* = 7-16/group (B-H). Box and whisker plots extend from 25th to 75th percentiles and show all points (B-H). Data were analyzed by two-way ANOVA with uncorrected Fisher’s Least Significant Difference (LSD) test (B-H). * *P* < 0.05; ** *P* < 0.01; *** *P* < 0.001; **** *P* < 0.0001

### THP enhances NETosis in human neutrophils with minimal impacts on cellular proteins

To determine if THP impact on NETosis extended to human models, we measured *in vitro* NETosis formation in primary human neutrophils with and without THP exposure. Peripheral circulating human neutrophils were isolated and incubated with THP purified from healthy human urine for 30 min at physiologic concentrations (50 µg/mL). After 2.5 h of stimulation with PMA, NETosis was measured by detection of fluorescently-labelled extracellular DNA as described previously(20). THP pretreatment increased levels of NETosis in PMA-stimulated cells but did not alter NETosis in unstimulated cells (**Fig. 5A**). To identify cellular processes impacted by the presence of THP, we performed tandem mass tag (TMT)-based quantitative proteomics of neutrophil cell pellets (*n* = 4 donors) under these same four conditions: mock-treated unstimulated (UnTx), THP-treated (THP), PMA-stimulated (PMA), and PMA-stimulated + THP-treated (PMA+THP). PMA stimulation was the primary driver of variation between samples as shown by PCA plot (**Fig. 5B**) and resulted in depleted neutrophil granule and NETs-related proteins likely due to the release of these proteins from activated cells during the 2.5 h incubation (**Supp.** Fig. 4A-B). Eight shared proteins were increased in THP (unstimulated) and PMA+THP (stimulated) groups compared to their mock-treated counterparts (**Fig. 5C, Supp. Table 2**). These proteins included THP itself (UMOD), and other known urinary proteins likely present in the purified THP preparation: apolipoprotein D, protein AMBP, kininogen, and galectin 3 binding protein (LGALS3BP)(70). The remaining three shared proteins were related to cellular metabolism (ACSS2, SLC16A9) and immune signaling (IL2RG). In unstimulated cells, 10 additional proteins (7 up and 3 down) were differentially abundant between THP-treated and mock-treated conditions (**Fig. 5D, Supp. Table 2**) and included several related to translational regulation and protein turnover (EIF2AK4, POLR3F, UBAC1), second messenger signaling (CD38), cytokine receptor signaling (RNF41), mitochondrial metabolism (GLDC, ALDH5A1), and phagosome acidification and fusion (RAB20), and chromatin remodeling (BICRAL). Gene ontology analyses identified mitochondrial respiratory chain complexes as significantly enriched in THP-treated conditions (**Fig. 5E**). In PMA-stimulated cells, 16 unique proteins (13 up and 3 down) were differentially abundant between THP-treated and mock-treated conditions (**Fig. 5F**, **Supp. Table 2**). These included proteins involved in second messenger and cell signaling (PDE7A, FCSK), transcriptional and translational regulatory proteins (PUM1, E2F3, ZFP36L2, GTPBP6, CCDC86), complement-related and immune related proteins (CD59, CXCL8, C4BPA, CTSW), intracellular trafficking and cytoskeleton arrangement (GIPC2, NCOA4, XIRP2), and DNA/chromatin remodeling proteins (DNASE1L1, BOD1). Gene ontology analyses identified tertiary granule and primary lysosome pathways as significantly enriched in THP-treated conditions and nonsignificant enrichment of chromosome condensation, autophagosome, and cytoskeleton pathways (**Fig. 5G**). Together, this proteomic profiling suggests THP induces subtle differential responses related to mitochondrial metabolism in the absence of PMA stimulation and impacts multiple nuclear, organelle, and cytoskeletal functions in stimulated conditions.

**Figure 5.**
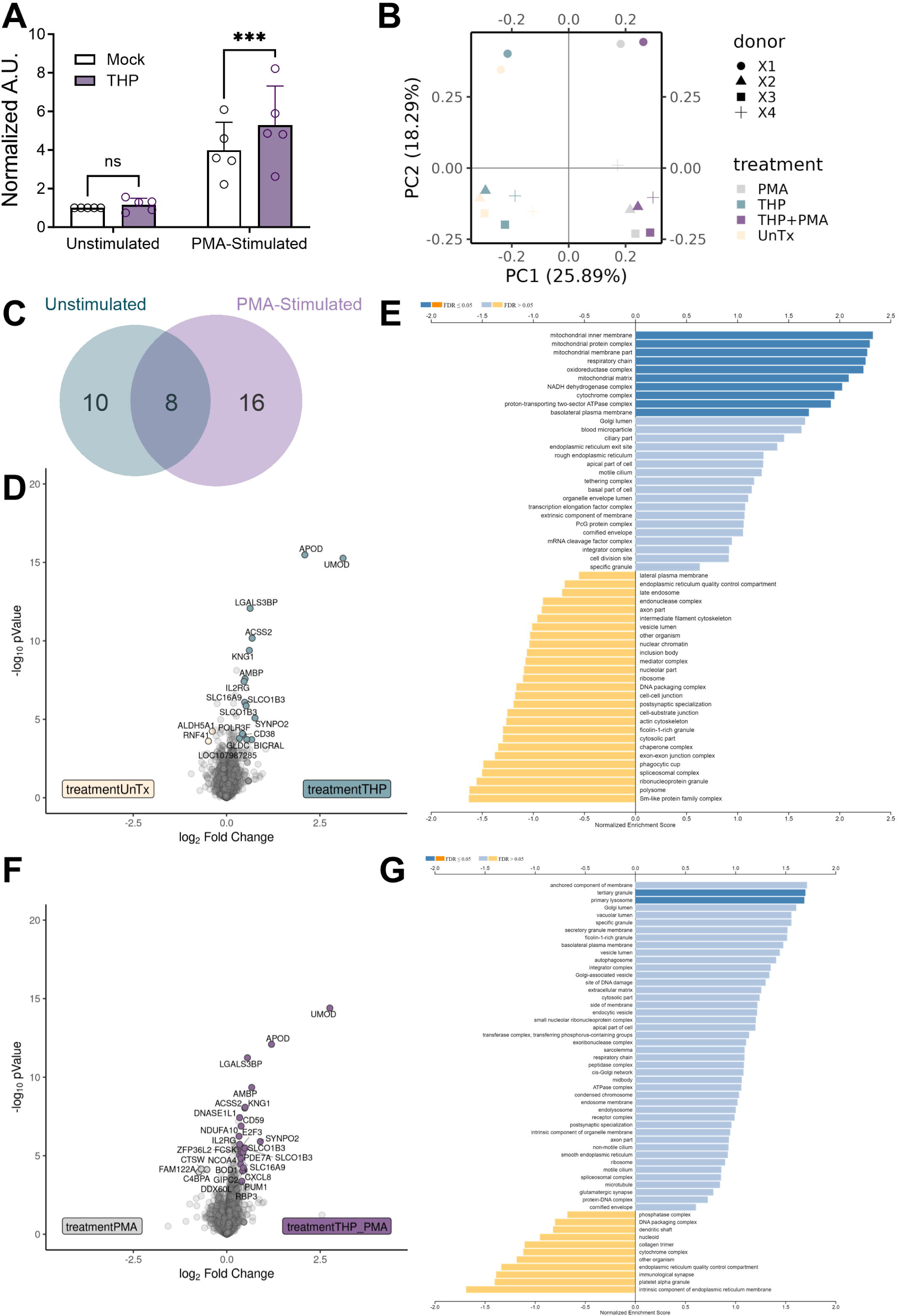
THP modestly alters neutrophil responses to NETosis stimulation by PMA. Peripheral human neutrophils were isolated, pretreated with THP, and stimulated phorbol 12-myristate 13-acetate (PMA) for 2.5-3 hours as described in Methods. (**A**) NET formation was assessed by released dsDNA (detected by PicoGreen dye) and expressed as arbitrary units (A.U.) of fluorescence normalized to mock-treated, unstimulated controls. Neutrophil cell pellets from four donors were subjected to tandem mass tag-based proteomics profiling. (**B**) Principal component analysis of neutrophils that were untreated (UnTx), treated with THP (THP), untreated with PMA stimulation (PMA), and THP-treated with PMA stimulation (THP+PMA). Each point represents an individual sample, colored by treatment, with paired donor samples indicated by matched symbol. (**C**) Venn diagram showing the proportion of proteins differentially detected in THP-treated samples compared to untreated samples in the presence (PMA and THP+PMA) vs. absence (UnTx and THP) of PMA stimulation. Volcano plot (**D**) and gene set enrichment analysis (**E**) of differentially identified proteins in untreated vs. THP-treated samples. NES = Normalized Enrichment Score. Volcano plot (**F**) and gene set enrichment analysis (**G**) of differentially identified proteins in PMA vs. THP+PMA samples. Experiments were performed as part of three independent experiments with data combined, *n* = 5 donors (A), or as part of one independent experiment, *n* = 4 donors (B-G). Box and whisker plots extend from 25th to 75th percentiles and show all points (A). Data (A) were analyzed by two-way ANOVA with Sidak’s multiple comparisons test. Differential proteins (C-I) were identified via Log_2_ fold change >1.25 and moderated t-test followed by multiple-hypothesis testing correction using the Benjamini– Hochberg procedure with a false discovery rate adjusted *P* <0.05. Individual proteins are listed in **Supplemental Table 2**. GSEA was performed with a gene set minimum of 10, a gene set maximum of 500, 2,000 permutations using the gene ontology cellular component gene sets.

### THP increases NETosis in human neutrophils in a ROS-dependent manner

To determine whether human neutrophil NETosis populations were similarly affected by THP as seen with murine neutrophils (**Fig. 4**), we modified our flow cytometry strategy for human neutrophils. Isolated peripheral human neutrophils were treated with purified human THP for 30 min and stimulated with PMA for 2.5 h as described above before staining and analysis via flow cytometry. Single cells were gated for the presence of extracellular/surface neutrophil granule content (MPO) and extracellular DNA (Sytox Orange) to identify double positive cells undergoing NETosis (MPO^+^SO^+^, P3) (**Fig. 6A**). P3 cells were further separated based on Hoechst brightness and Live/Dead staining into nonclassical NETosis (Hoechst^lo^ Live/Dead^−^, P4) and classical NETosis (Hoechst^hi^ Live/Dead^+^, P5) subsets (**Fig. 6A**). Consistent with the fluorescence-based NETosis assay (**Fig. 5A**), THP treatment significantly increased total NETosis in PMA-stimulated conditions, but not in the unstimulated cells (**Fig. 6B**). Furthermore, THP treatment enhanced both nonclassical (**Fig. 6C**) and classical NETosis (**Fig. 6D**) subsets specifically in PMA-stimulated conditions. Classical and nonclassical NETosis are dependent on NADPH oxidase 2-mediated production of reactive oxygen species(31, 71, 72) and hydrogen peroxide (H_2_O_2_) as an exogenous source of ROS is sufficient to stimulate NETosis *in vitro*(28). To examine the importance of ROS on THP-mediated effects of NETosis, we compared nonclassical and classical NETosis subsets in the presence of PMA, H_2_O_2_, or PMA and a NADPH oxidase inhibitor diphenyleneiodonium (DPI). The THP-mediated increase in nonclassical NETosis occurred in the presence of both PMA and H_2_O_2_ but was abrogated with the addition of DPI (**Fig. 6E**). In contrast, no significant differences were observed in classical NETosis subsets under these same conditions (**Fig. 6F**). Together, these data suggest that THP-mediated effects are in part dependent on ROS, specifically in cells undergoing nonclassical NETosis.

**Figure 6.**
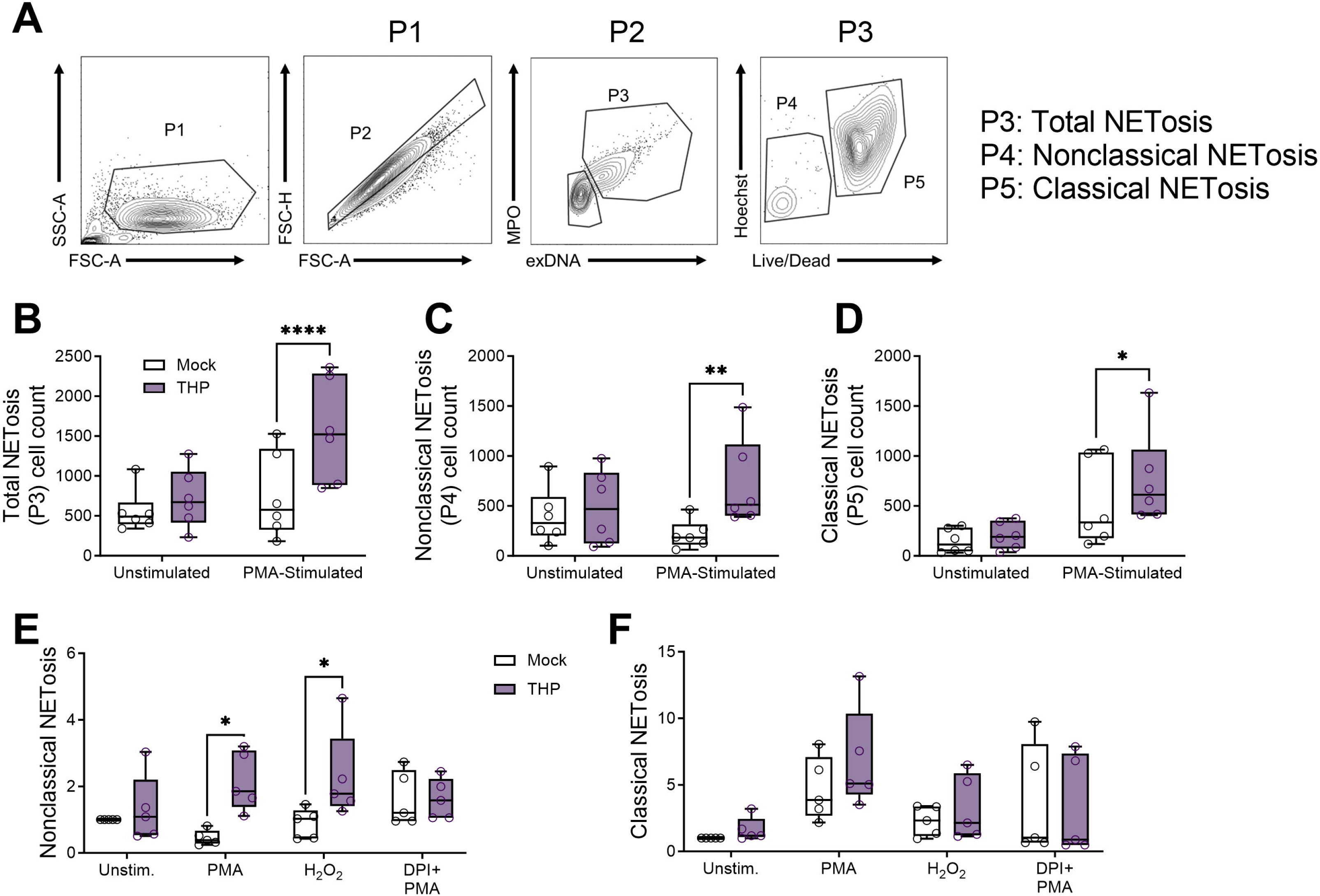
THP exposure increases NETosis in human neutrophils. Peripheral human neutrophils were isolated, pretreated with THP, and stimulated phorbol 12-myristate 13-acetate (PMA) for 2.5 hours. (**A**) Gating strategy for quantifying neutrophil NETosis (extracellular DNA [Sytox Orange]+, MPO+, P3) subpopulations of interest with a focus on nonclassical NETosis (Hoechst^var^, Live/Dead-) and classical NETosis (Hoechst^hi^, Live/Dead+). (**B**) Total NETosis (P3) cell counts across treatment groups. (**C**) Nonclassical NETosis (P4) cell counts across treatment groups. (**D**) Classical NETosis (P5) cell counts across treatment groups. Peripheral human neutrophils were pretreated with THP or mock-treated as above and subsequently stimulated with either PMA, H_2_O_2_, or PMA + DPI (diphenyleneiodonium, an inhibitor of ROS). Frequency of nonclassical NETosis (P4) cells (**E**) or classical NETosis (P5) cells (**F**) normalized to frequency of unstimulated cells from the same donor. Experiments were performed in at least four independent experiments with data combined, *n* = 6 donors (B-D), or *n* = 5 donors (E-F). Box and whisker plots extend from 25th to 75th percentiles and show all points (B-F). Data were analyzed by two-way ANOVA with Sidak’s multiple comparisons test (B-F). * *P* < 0.05; ** *P* < 0.01; **** *P* < 0.0001.

### THP alters cell shape and chromatin staining during NETosis as measured by imaging flow cytometry

Prior to DNA release, cells destined for NETosis undergo multiple cellular remodeling events including cytoskeletal and endoplasmic reticulum disassembly(73), vacuolization, autophagy, and superoxide production(74), and lastly, chromatin swelling and nuclear envelope rupture(75). Live cell imaging or imaging flow cytometry techniques combined with mathematical modeling and/or machine learning have revealed predictable morphologic changes that can delineate NETosis from other forms of cell activation and death(32, 64, 73, 76, 77). To assess whether THP altered neutrophil morphology during NETosis, we adapted an imaging flow cytometry method and algorithm from prior work(76) to identify NETs, NET precursors, and other forms of cell death. Using this method, cells can be distinguished into six cell types: healthy (Type I), live cell decondensed nuclei (Type II), NETs (Type III), DNA fragments (Type IV), dead cell condensed nuclei (Type V), and dead cell diffuse nuclei (Type VI). Peripheral human neutrophils were pretreated with THP or an estimated equivalent amount of sialic acid (Sia, 500 ng/mL) and stimulated with PMA for 2.5 hours. Cells were stained with α-MPO-FITC, Sytox Orange, Hoechst 33342, and Live/Dead Near I/R, subjected to imaging flow cytometry, and gated as shown in **Fig. 7A**. Cells were separated from debris based on brightfield (BF) area and Hoechst^+^ staining. NETs (Type III) and DNA NET fragments (Type IV) were distinguished by higher extracellular DNA (calculated by the SO staining beyond the BF margins of the cell) area and higher or lower Hoechst intensity respectively. The remaining cells were further gated to collect focused, single cell images and separated based on SO intensity (indicating membrane permeability) and Hoechst area (indicating nuclear area). Dead cells with condensed nuclei (Type V) and dead cells with diffuse nuclei (Type VI) were marked by higher SO intensity and delineated by lower and higher Hoechst area respectively. Healthy cells (Type I) and live cells with decondense nuclei were demarcated by Hoechst area. Representative images of each cell type are shown in **Fig. 7B** with dead cell types (V and VI) confirmed by staining Live/Dead^+^ while all other types were Live/Dead^−^. No differences in Hoechst+ populations were observed across groups (**Supp.** Fig. 5A). Although individually, NETs and NET fragment frequencies did not differ between groups (**Supp.** Fig. 5B-C), the sum of NETs and NET fragment frequencies were significantly higher in the PMA + THP group compared to mock-treated controls (**Fig. 7C**). Additionally, the PMA + THP group exhibited higher frequencies of Type II (**Fig. 7D**) and decreased Type V frequencies (**Fig. 7E**) compared to mock-treated controls. Other subsets (Type VI and Type I) were not significantly different between groups (**Fig. 7F-G**). Using the Feature Finder Analysis tool in the IDEAS 6.3 software, we also identified the circularity feature, which gives higher scores to features closely resembling a circle, as significantly higher in both PMA + THP and PMA + Sia groups compared to mock-treated controls (**Supp.** Fig. 5D). Overall, these analyses reveal that THP, in the presence of a NETosis stimuli, enhances the frequency of NETs (Type III), NETs fragments (Type IV), and NETs precursors (Type II) over baseline conditions.

**Figure 7.**
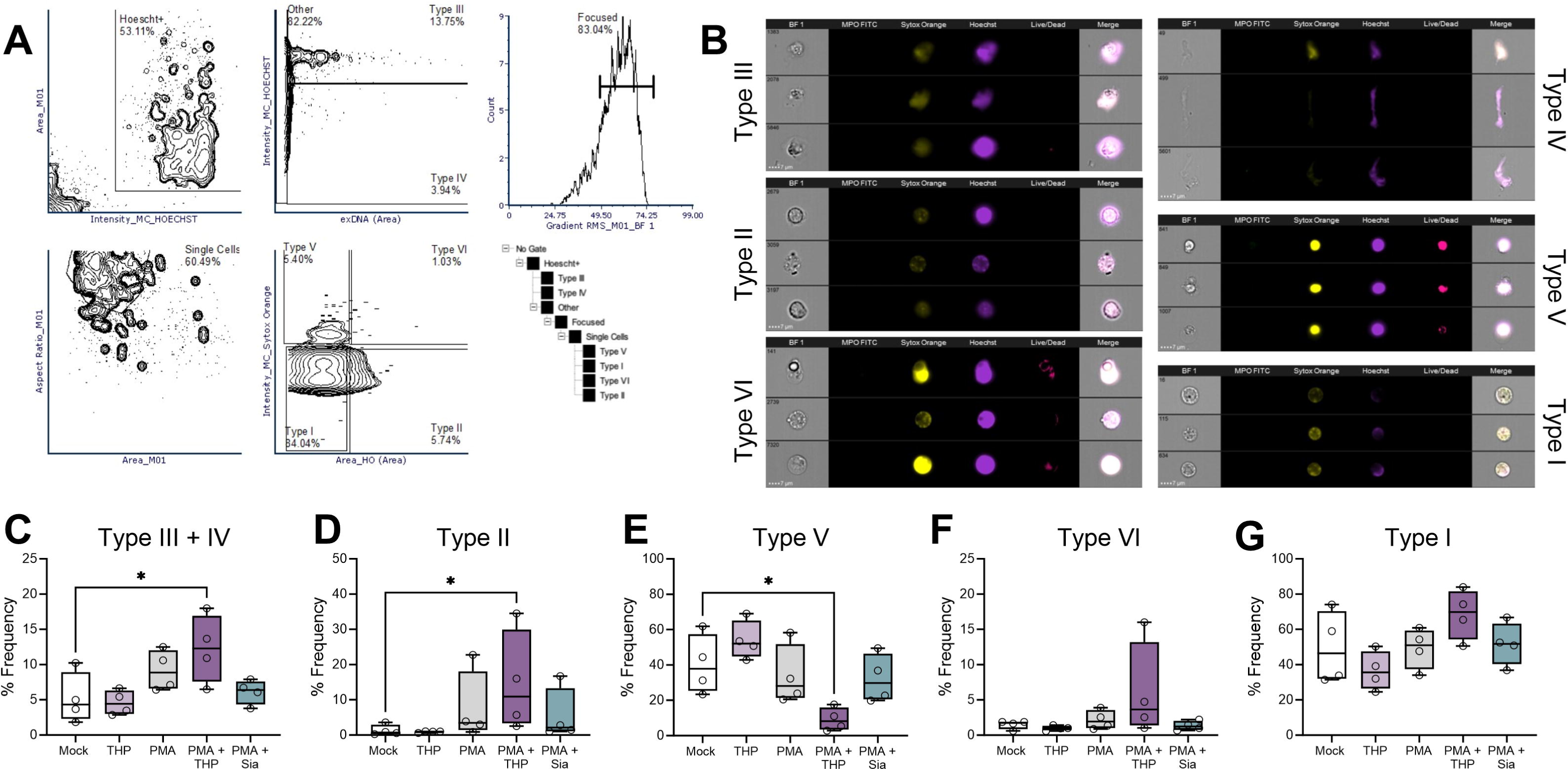
THP exposure alters proportions of NETs and other cellular morphologies as determined by imaging flow cytometry. Peripheral human neutrophils were isolated, pretreated with THP or sialic acid, and stimulated phorbol 12-myristate 13-acetate (PMA) for 2.5 hours. Cells were stained with anti-MPO FITC, Sytox Orange (non-membrane permeable nucleic acid dye), Hoechst (membrane-permeable nucleic acid dye), and Live/Dead stain (non-membrane permeable amine-reactive dye) and visualized for fluorescence and brightfield (BF) images on an imaging flow cytometer. (**A**) Gating strategy of human neutrophils subpopulations with representative images shown (**B**). NETs (Type III) are gated from Hoechst+ cells based on high Hoechst intensity and extracellular DNA area (Sytox Orange staining beyond cell margins), whereas NET DNA fragments (Type IV) were gated as high extracellular DNA with lower Hoechst intensity. Remaining cell populations are collected from focused cells and gated based on Sytox Orange intensity (indicating cell permeability) and Hoechst area to monitor nuclear morphology. Two dead cell populations (high Sytox Orange intensity and confirmed by Live/Dead staining) were separated into dead cells with condensed nuclei (Type V) and dead cells with decondensed nuclei (Type VI). Live cell populations (low Sytox Orange intensity and absence of Live/Dead stain) were separated into cells with decondensed nuclei (Type II) and live cells (Type I). Frequency of NETs and NETs DNA fragments (Type III and IV) (**C**), live cells with decondensed nuclei (Type II) (**D**), dead cells with condensed nuclei (Type V) (**E**), dead cells with decondensed nuclei (Type VI) (**F**), and live cells (Type I) (**G**). Experiments were performed in four independent experiments with data combined, *n* = 4 donors. Box and whisker plots extend from 25th to 75th percentiles and show all points (C-G). Data were analyzed by one-way ANOVA with Holm-Sidak’s multiple comparisons test (C-G). * *P* < 0.05.

## DISCUSSION

Despite abundant evidence supporting the critical independent roles of THP and neutrophils in protecting against UTI, few studies have investigated direct interactions between these two host defenses. In this work, we build upon recent findings of THP regulation of neutrophil function and provide a more detailed characterization of the histopathological and immunological consequences of THP deficiency in the urinary tract during UTI. From our *in vivo* experiments, neutrophils emerged as a key cell type; THP-deficient mice displayed altered neutrophil proportions and NETosis during UTI, combined with improved bacterial control upon neutrophil depletion. In prior work, we demonstrated that THP’s terminal sialic acids binding neutrophil Siglec-9, inhibiting ROS production among other regulatory effects(50). Presently, employing multiple methods including imaging flow cytometry, we show that in the presence of NETosis stimuli, THP enhances characteristics associated with NETosis such as nuclear decondensation. We propose that this function, coupled with THP-mediated dampening of excessive neutrophil activation(50), is integral to protecting the urinary tract from infectious and inflammatory insults.

Since their initial reports in 2004(39, 40), two independent transgenic THP KO mouse lines have consistently demonstrated that THP deficiency increases susceptibility to urinary pathogens(39–42, 78) and aggravates renal pathologies(37, 79, 80). Although histologically similar at baseline, THP KO mice display more severe renal necrosis and neutrophil infiltration upon acute kidney injury(52). Additionally, they show increased bladder lamina propria thickness and neutrophil invasion of the uroepithelium during *Klebsiella pneumoniae* or *Staphylococcus saprophyticus* UTI(78). Our findings build upon these prior studies, revealing that THP KO mice exhibit more severe histopathological alterations in both bladder and renal tissues during acute UPEC UTI.

In both THP KO lines, increased neutrophil recruitment to affected tissues is a primary phenotype during urinary tract injury or exposure to inflammatory stimuli(50, 79). Likewise, in this study we identified neutrophils as the predominant immune cell population impacted by THP deficiency. In contrast to a prior study using the other THP KO transgenic line(53), we did not observe elevated renal neutrophils in our THP KO mice at baseline; this divergent phenotype may be due to differences between *Neo* cassette placement between the two independent THP KO transgenic lines or variations in the methods used to evaluate neutrophil populations. The heightened tissue damage in THP KO mice may, in part, be driven by excessive neutrophil activation as blocking neutrophil recruitment through chemokine depletion ameliorates tissue damage in THP KO mice(79). While neutrophil depletion worsened bacterial burdens in the bladders of WT mice, their depletion in THP KO mice resulted in reduced bladder burdens. This highlights the importance of neutrophils in both bacterial killing and tissue damage and is supported by a prior study demonstrating that high levels of neutrophil depletion (200 μg/dose) worsened bacterial burdens and promoted chronic infection, while partial neutrophil depletion (10 μg/dose) lessened the incidence of chronic infection(17).

Human THP possesses eight N-glycosylation sites with high-mannose and bi-, tri-, or tetra-antennary complex types(46) and these glycans are crucial to THP structure and function. THP glycosylation facilitates direct interactions with both neutrophils and *E. coli*(43, 50). Altered N-glycan profiles, including reduced galactose and α2–6-linked sialic acid, have been reported in UTI patients compared to healthy controls(81). To our knowledge, this work represents the first report of N-glycosylation patterns on murine THP. Despite being only 70% identical at the amino acid sequence level, mouse THP possesses the same N-glycan sites as human THP(82), and here we show that the glycan structures themselves are also similar including the shared, abundant tetra-antennary, tetra-sialylated and fucosylated N-glycan. Reduced sialic acid levels have been reported in THP from patients with UTI, interstitial cystitis, and kidney stones(61, 62, 81); however, we did not observe any changes in THP total sialic acid levels during murine UTI. Due to low urine volumes, we pooled from multiple mice and collected over the first 72 hours following infection. Thus, it is possible that reduction in sialic acid occurs at later periods during UTI. Additionally, we determined that the primary sialic acid modification on murine THP is Neu5Ac rather than Neu5Gc, a common mammalian sialic acid not present in humans(83). Together, these data demonstrate that THP sialic acid-neutrophil Siglec signaling remains intact during UTI and further highlight the utility of mouse models in studying mechanistic functions of THP glycans in the urinary tract.

This study provides the first visualization of NETs during murine UTI (**Fig. 4**), complementing several *in vivo* studies that provide evidence of the importance of NETosis in UTI. Both *Irf3*^-/-^ and *Ifn*β^-/-^ mice present with abscess formation and tissue damage during UTI indicating defective neutrophil responses(84). This may involve reduced NET formation as the type I IFN response(85), and specifically IFN-β(86), drive NETosis in mouse models of lung infection. A recent study using protein-arginine deiminase type 4 (PAD4) KO mice as a model for reduced NETosis formation found that PAD4 KO mice displayed higher bladder and kidney bacterial burdens in UPEC UTI(87). Proteomic studies of urine from UTI patients have identified NET-associated proteins in samples from bacterial and fungal infections, suggesting that NETosis may be a conserved host urinary defense against a wide range of uropathogens(34). Additionally, neutrophil NETosis in response to UPEC UTI was demonstrated on a bladder-on-a-chip model using diluted human urine as the luminal medium; thus, THP would be present in this system(36). These *in vitro* studies did not distinguish subtypes of NETosis and did not associate NET formation with outcomes. However, future work using these or similar platforms could determine the contribution of THP to neutrophil migration, NETosis, and resolution of infection in a dynamic model of the uroepithelium.

In this study, we used proteomics to parse the signaling pathways impacted by THP treatment in the presence and absence of PMA stimulation. Our findings were generally in line with other proteomic-based studies of PMA-induced NETs in human(88) and mouse(89) neutrophils, with some overlap in cellular responses to platelet-activating factor, another stimulus of NETosis(90). PMA stimulation resulted in organelle and cytoskeleton-related proteins which may reflect cytoplasmic changes occurring prior to NETosis(73, 75) or during the early, active stages of NETosis(75). In this study, we found that THP itself minimally altered protein profiles, with modest increases in tertiary granule and primary lysosome pathways in the presence of PMA. Neutrophil retention of granules is thought to contribute to membrane breakdown during NETosis(91), and autophagy is required for intracellular chromatin decondensation(74), suggesting multiple effects of THP exposure. However, there are several limitations impacting the interpretation of this data set. It is possible that measuring changes in relative abundances of protein levels is not the most suitable method to evaluate NETosis induction. Protein translation is dispensable for NOX-dependent or NOX-independent NETosis although transcriptional changes are observed as soon as 30 minutes post-exposure to stimuli(92). Another limitation of our approach is that we evaluated proteomics of cell pellets that remained after 2.5 h of stimulation, thus observing lower levels of key markers of NETosis and neutrophil degranulation that were released from activated and/or lysed cells. Additionally, since only one time point was evaluated, differences in protein kinetics were not captured. Even so, by comparing PMA-stimulated cells in the presence or absence of THP, we identified multiple differentially abundant proteins linked to cytoplasmic and chromatin remodeling and these candidates are the focus of future studies.

Both the classical (suicidal) and mitochrondrial types of NETosis are dependent on NADPH oxidase 2 (NOX2) activity(31, 71, 72, 93). Neutrophils deficient in NADPH oxidase fail to induce actin and tubulin polymerization and NET formation upon stimulation(94). Nonclassical (vital) NETosis, where the cell membrane initially remains intact, may be NOX-independent at early time points (< 1h) but becomes NOX-dependent at later time points(33). Nonclassical NETosis is characterized by more extensive histone citrullination, delayed ROS release, dilatation of the nuclear envelope prior to rupture, and the presence of extracellular DNA NETs despite having intact plasma membranes(33, 95). We observed that THP enhanced nonclassical NETosis, and to a lesser extent, classical forms, in a manner dependent on ROS *in vitro* (**Fig. 6**). Additionally, a reduced frequency of apoptotic cells was observed in PMA-stimulated THP-treated neutrophils (**Fig. 7**). In prior work, Siglec 9 crosslinking reduced apoptosis, but promoted non-apoptotic cell death, in GM-CSF-stimulated neutrophils in a ROS-dependent manner(96). It is interesting to speculate that THP, through Siglec-9 mediated inhibition of apoptosis, may allow more opportunity for stimulated cells to undergo NETosis. This would also explain the observed increased proportion of cells with dilated nuclear envelops and decondensed chromatin in the presence of intact plasma membranes. A limitation of these *in vitro* assays is that they were not performed in the context of human urine or infection; thus, cellular activation may differ. Nonetheless, we still observed similarities in THP-associated increases in nonclassical NETosis in both mouse UTI and human neutrophils *in vitro* using parallel methodologies.

Imaging flow cytometry has been used in various studies to categorize NETosis. In one study, the categorization of ‘suicidal’ and ‘vital’ NETosis based on neutrophil morphology in response to LPS stimulation(97). In this work, Zhao *et al* observed populations with diffuse MPO and nuclear (Hoechst) staining, indicating decondensed nuclei, which they termed suicidal NETosis, and another population that were elongated with large blebs at one pole and nuclear and granular contents at the other pole, which they termed vital NETosis, hypothesizing that the nuclear material was being extruded leaving anuclear cells with intact membranes behind. However, we did not observe this morphology in our assays, possibly due to differences in time course (1 h vs. 2.5 h) and stimuli (LPS vs. PMA). Interestingly, they also described both suicidal and vital NETosis occurring after 4 h of PMA stimulation using the same morphologic characterization(97). Another study used nuclear morphology (normal or decondensed) and histone H4 citrullination to assess NETosis in response to hemin, PMA, LPS, and IL-8 over a 1 h treatment period(77). To differentiate between NETosis and other forms of cell death, another study used a combination of cell permeable and non-permeable nucleic acid dyes and cell boundaries defined by bright field images(76). We adapted this method to our samples and found striking similarities in cell morphologies with the prior work(76) even with differences in staining (e.g. MPO vs. NE, Sytox Orange vs. Sytox Green, inclusion of Live/Dead viability dye). Observed differences in cell type frequencies were likely different due to differences in experimental methods (e.g. we did not use Percoll). Our results further validate this methodology as a robust pipeline for rapidly distinguishing NETs from other forms of cell death and we recommend the addition of a Live/Dead stain to this pipeline to confirm cell membrane permeability. It is possible additional cell types could be distinguished from this data set: for instance, while some cells undergoing NETosis stained with MPO indicating degranulation, others did not (**Supp.** Fig. 5E). We found that sialic acid alone did not alter cell morphologies to the extent that THP did in PMA-stimulated cells with the exception of increased cell circularity. This suggests that endogenous proteins such as THP may differentially signal through Siglec-9 compared to free sialic acid or pathogen-mediated engagement of Siglec-9(98, 99) to alter NETosis.

In summary, our study reveals that THP modulates neutrophil NETosis in both animal models and human neutrophils *in vitro*. We postulate that this activity provides an additional layer of THP-mediated protection against UTI. Acting as a multi-faceted host defense through both blocking pathogen adherence and modulating immune cell function, pharmacologic manipulation of THP(100, 101) may emerge as a promising therapeutic target to improve outcomes and prevent UTI in susceptible populations.

## METHODS

### Sex as a biological variable

This study exclusively examined female mice. It is unknown whether the findings are relevant for male mice. Both male and female human donors were included; however, due to small sample sizes, we are underpowered to determine sex-dimorphic effects in this study.

### Bacterial strains and growth conditions

Wildtype uropathogenic *E. coli* strain UTI89(102) was grown overnight at 37°C in Luria-Bertani (LB) broth with shaking. Stationary phase overnight cultures were then centrifuged for 5 minutes at 3,200 × *g* and resuspended in an equivalent volume of PBS.

### Murine model of UPEC urinary tract infection

Wild type (WT) THP^+/+^ and THP^-/-^ (THP KO) mice(39) were bred and maintained at UCSD and BCM. Groups were randomly assigned and mice were housed 4-5 animals per cage. Mice ate and drank ad libitum. All animals used in this study were females aged 2 to 5 months.

UPEC strain UTI89 was prepared as described above. Mice were anesthetized with inhaled isoflurane and infected via transurethral inoculation with approximately 10^8^ CFU in 50 μl of PBS as described previously(54). Twenty-four hours to ten days post-infection, mouse urine was expressed and/or bladders and kidneys were collected. Tissues were homogenized in tubes containing 1.0-mm-diameter beads (Biospec Products; catalog number 11079110z) using a MagNA Lyser instrument (Roche Diagnostics). Serial dilutions of homogenized organs were plated on LB agar and enumerated the following day. Urine samples were either plated for CFU on LB agar or processed for flow cytometry or microscopy as described below. For partial neutrophil depletion, mice were administered 10 µg of anti-Ly6G (clone 1A8, catalog no. BE0075-1; BioXCell) or rat IgG2a isotype control (clone 2A3, catalog no. BE0089; BioXCell) in 100 µL of sterile PBS i.p. just prior to bacterial inoculation. Mice received additional antibody injections on day 2 and 4 post-inoculation. For diclofenac treatment, mice were administered diclofenac sodium salt (Thermo Scientific Chemicals) at a targeted 30 mg/kg/day dose as described previously(56). To achieve this dosage, we determined that the average mouse weighed 20 g and consumed 3 mL of water daily thus mice were given diclofenac sodium salt (0.2mg/mL) in drinking water on days 0 through 6 post-inoculation.

### Tissue pathology assessment and scoring

Bladder and kidney tissues were collected at day 1 and 3 post-inoculation, fixed in 10% neutral buffered formalin for 24 h, and dehydrated by an ethanol gradient and embedded in paraffin. Tissue sectioning (4 µm) and hematoxylin and eosin (H&E) staining was performed by the UC San Diego Comparative Phenotyping Core. Tissue sections were examined by a veterinary pathologist blinded for treatment (UPEC versus mock-infection) and genetic background (WT or THP KO). Severity of bladder inflammation was scored based on number of infiltrating cells, degree of tissue damage, and by the presence or absence of visible bacteria. Scores ranged from 0 (no lesions) to 4 (severe lesions). Minimal to mild lesions (0–2) consisted of small numbers of infiltrating inflammatory cells and intraluminal bacteria/debris. Severe lesions (3–4) involved fibrinoid necrosis of submucosal blood vessels, submucosal edema, and micropustule formation within the urothelium. Brightfield images were collected using an Echo Revolve microscope at 400X (bladders) and 200X (kidneys) magnification.

### Flow cytometry of bladder and kidneys

Bladder and kidney tissues were subjected to flow cytometry as adapted from prior work(19). Tissues were finely minced and incubated in RPMI 1640 containing 4 mg/mL collagenase and 50 U/mL DNase for 1 h at 37°C with manual pipetting every 15 min. Samples were passed through 40-μm filters and washed in RPMI 1640 medium with 10% FBS. Kidney samples were subjected to RBC lysis by cells in 0.2% (w/v) saline for 30 seconds with gentle mixing and then stopping lysis by adding an equal volume of 1.6% (w/v) saline. Cells were blocked with 1:200 mouse BD Fc-block (BD Biosciences) for 15 min on ice in PBS with 1 mM EDTA, 1% FBS, and 0.1% sodium azide. Cells were stained for 30 min on ice using the following antibodies (all at 5 μg/mL): anti-CD11b-FITC (clone M1/70, catalog no. 553310; BD Biosciences), anti-CD11c-PerCP-Cy5.5 (clone N418, catalog no. 45-0114-82; eBioscience) or anti-CD11c-BV786 (clone N418, catalog no. 117335; BioLegend), anti-Ly6G-APC (clone 1A8, catalog no. 127614; BioLegend), anti-MHC-II-APC-Fire750 (clone M5/114.15.2, catalog no. 107652; BioLegend) or anti-MHC-II-BV650 (clone M5/114.15.2, catalog no. 563415; BD Biosciences), and anti-CD45-BV510 (clone 30-F11, catalog no. 103138; BioLegend) or anti-CD45-BV605 (clone 30-F11, catalog no. 563053; BD Pharmingen). Samples were washed 1X, resuspended in fresh FACS buffer, and run on a BD FACSCanto II (BD Biosciences). Samples were gated on unstained cells as described in **Fig. 2A** and positive signals were determined using single-stain controls, and data were analyzed with FlowJo version 10.9.0 (FlowJo LLC).

### Urine Sediment Scoring

Urine sediment scoring was performed as described previously(17). Mouse urine was diluted 1:10 and centrifuged onto glass slides using a Cytospin 3 (Thermo Shandon) at 1000 rpm for 3 minutes. Slides were covered with Wright-Giemsa stain for 30 seconds, washed twice with water, and visualize by light microscopy on Olympus BX41 brightfield microscope at 200X magnification (hpf). The average number of polymorphonuclear (PMN) cells was calculated from counting 2 independent fields and scored by a semi-quantitative scoring system of 0-5: 0, < 1 PMN/hpf; 1, 1–5 PMN/hpf; 2, 6–10 PMN/hpf; 3, 11–20 PMN/hpf, 4, 21–40 PMN/hpf, and 5, > 40 PMN/hpf.

### Flow cytometry of murine urine

Mouse urine was subject to flow cytometry analyses as described previously(50) with several modifications. Urine volume was recorded and then passed through a 40-μm filter. Cells were washed in PBS and resuspended in 50 μL of FACS buffer (1mM EDTA, 1% FBS, 0.1% sodium azide in PBS). The following antibodies (0.5 μg/mL) and dyes (concentrations provided below) were added: Anti-CD11b-FITC (clone M1/70, catalog no. 553310; BD Biosciences), anti-Ly6G-APC (clone 1A8, catalog no. 127614; BioLegend), Live/Dead Near IR (1:200 of stock, catalog no. L34975; Thermo Fisher Scientific), Sytox orange (100 nM, catalog no. S34861; Thermo Fisher Scientific), and Hoechst 33342 (200 nM, catalog no. 62249; Thermo Fisher Scientific). After a 30-minute incubation on ice, samples were washed 1X, resuspended in fresh FACS buffer, and run on a BD FACSCanto II (BD Biosciences). Samples were gated on unstained cells as described in **Fig. 4A** and positive signals were determined using single-stain controls, and data were analyzed with FlowJo version 10.9.0 (FlowJo LLC).

### Immunofluorescence of murine urine

Mouse urine, collected 24 h post-infection, was diluted 1:20 and centrifuged onto glass slides using a Cytospin 3 (Thermo Shandon) at 1000 rpm for 5 minutes. Samples were fixed in 1% paraformaldehyde for 10 min, washed 1X, and permeabilized with 0.1% TritonX at room temperature. Cells were washed in PBS + 0.01% Tween20 (PBST) and blocked for 1 h in PBST with 10% horse serum and 1% BSA. Cells were incubated overnight at 4°C with primary antibodies: sheep anti-THP polyclonal antibody (1:40, catalog no. AF5175; R&D Systems), goat anti-MPO polyclonal antibody (1:200, catalog no. AF3667; R&D Systems), and rabbit anti-Histone H3 (citrulline R2 + R8 + R17) polyclonal antibody (1:100, catalog no. ab5103; abcam) diluted in PBST + 1% BSA. The anti-THP antibody was conjugated with FITC using the FITC Conjugation Kit Lightning-Link (abcam ab102884) kit per manufacturers’ instructions. Cells were washed 2X in PBST and incubated for 1 h at room temperature with secondary antibodies anti-Goat IgG-AF647 (1:250, catalog no. A-21469; Thermo Fisher Scientific) and anti-rabbit IgG-Texas Red (1:400, catalog no. ab6800, abcam) to visualize MPO and H3cit respectively, and nuclei were stained with NucBlue Fixed Cell ReadyProbes Reagent (catalog no. R37606, Thermo Fisher Scientific). Slides were mounted, cured, and fluorescence images collected using an Echo Revolve microscope at 600X magnification using filter configurations for DAPI, FITC, TxRED and CY5.

### Tamm-Horsfall glycoprotein quantification and purification

Mouse urine was expressed every 3-4 hours up to 3 times a day for the first 96 hours post-inoculation from WT and THP KO mice and mock-infected controls. Total THP levels of a subset of urine samples (diluted 1:10) was determined by ELISA (catalog no. DY5144-05, R&D Systems) per manufacturers’ instructions. WT samples falling below the limit of detection were excluded from analyses. For murine THP purification, urine was pooled from multiple mice and time points based on genotype (WT vs. THP KO) and infection status (UPEC-infected vs. mock). For human THP purification, urine from healthy male and female donors was collected and stored at 4°C until THP purification. THP was purified via an adapted protocol(62). Briefly, 2.5 g of diatomaceous earth (DE) was combined with Milli-Q water to create a 50 mL slurry. The DE slurry was passed through a 60 mL Büchner funnel with vacuum filtration to create a DE layer. The DE layer was washed with 50 mL of Milli-Q water followed by 50 mL of PBS. Urine (approximately 10 mL) was filtered through the DE layer, followed by 50 mL of PBS, and the DE layer was dried under vacuum for 2 min. The DE layer was transferred to a 50 mL conical and THP bound to the layer was solubilized by adding Milli-Q water (50 mL) under gentle rocking for 30 min. Samples were centrifuged at 3220 × *g* for 30 min and supernatant containing THP was run through an Amicon® Ultra 50 kDa filter at 3220 × *g* for 15 min in multiple batches. Filters were washed 3X with 15 mL of Milli-Q water. Total protein in retentate was measured using BCA assay (Pierce, Catalog no. 23225) per manufacturers’ instructions, lyophilized, and stored at –80°. Purity of both mouse and human was confirmed by running purified THP on a 4-12% Bis-Tris polyacrylamide gel and staining with SimplyBlue™ SafeStain (Thermo Fisher Scientific). THP was visualized as a single band at ∼ 85 kDa (**Supp.** Fig. 6).

### THP glycan analyses

For measurement of sialic acid, purified mouse THP samples (25 µg of protein) were hydrolyzed using 2M acetic acid at 80°C for 3 h to release sialic acids (Neu5Ac and Neu5Gc). Excess acid was removed via evaporation using a speed vacuum and hydrolyzed sialic acids were then dissolved in known volume of ultrapure water and tagged with DMB reagent at 50°C for 2.5 h. Finally, DMB-tagged sialic acids (2 µg dissolved in ultra-pure water) were injected on a Reverse Phase Ultra Pressure Liquid Chromatography Florescence Detector (RP-UPLC-FL, Waters Acquity UPLC) set at λex 373 nm and λem 448 nm on a Acquity UPLC BEH C18 1.7µm, 2.1mm x 50mm column (Waters, cat. No. 186002350). Solvents included 7% methanol with 0.1% TFA in HPLC water and acetonitrile with 0.1% TFA and the flow rate was set to 0.4 mL/min. Sialic acids were quantified by comparing with known amount of standard mixture (Neu5Gc and Neu5Ac) purchased from Sigma-Aldrich.

N-linked glycans were enzymatically released from purified mouse THP using PNGase-F kit (catalog no P0709S, New England Biolabs). N-glycans were then purified from the reaction mixture containing denaturing buffer and de-N-glycosylated proteins by solid phase extraction method using Sep-Pak C18 (1 cc Vac-cartridges, Waters) and poly-graphitized charcoal cartridge (Supelclean Envi-Carb, Supelco). Purified N-glycans were per-methylated and analyzed by MALDI-Tof/Tof mass spectrometry (Bruker, AutoFlex) in positive mode. Briefly, N-glycans were dried completely and re-dissolved in anhydrous DMSO, followed by per-methylation using NaOH slurry in anhydrous DMSO and CH3I. Per-methylated glycans were extracted with chloroform, and dried completely using dry nitrogen flush. The permethylated N-glycans were dissolved in mass spec grade MeOH and mixed in 1:1 (v/v) ratio with Super-DHB (MALDI matrix) before spotting. Sample (1µL) were spotted and allowed to air dry before acquiring mass spectra. All MALDI mass spectral data on permethylated N-glycans were acquired in positive and reflectron mode. Finally, the mass spectral data was analyzed and plausible N-glycan structures were annotated using GlycoWork Bench software selecting CFG database.

### Human neutrophil isolation and NETosis assays

Under approval from UC San Diego IRB/HRPP and BCM IRB, venous blood was obtained after informed consent from healthy adult volunteers, with heparin as an anticoagulant. Neutrophils were isolated using PolymorphPrepTM (Axis-Shield) to create a density gradient by centrifugation according to the manufacturer’s instructions. Fluorescence-based quantification of neutrophil extracellular traps (NETs) was performed as described previously(20). Briefly, isolated neutrophils were plated on 96-well tissue culture plates at 2 × 10^5^ cells/well. Cells were pretreated with 50 μg/mL of purified human THP or mock-treated, and incubated at 37°C in 5% CO_2_ for 30 min, and then incubated for an additional 3 h with phorbol 12-myristate 13-acetate (PMA, Sigma Aldrich) PMA (25 nM) to induce NET production(20). Micrococcal nuclease was then added at a final concentration of 500 mU/mL for 10 min to digest extracellular DNA. Plates were centrifuged at 200 g for 8 min; sample supernatant was then collected and transferred to a new 96-well plate. DNA was quantified using a Quant-iT PicoGreen® dsDNA Assay Kit from Life Technologies (Carlsbad, CA, USA), with fluorescence detected on intensity (485 nm excitation and 530 nm emission) measured by an EnSpire Alpha Multimode Plate Reader (PerkinElmer).

### Human neutrophil proteomics and analyses

Peripheral human neutrophils were isolated as above and pretreated with 50 μg/mL of purified human THP or mock-treated, and incubated at 37°C in 5% CO2 for 30 min, and then incubated for an additional 2.5 h with PMA (25 nM). Cells were pelleted by centrifugation at 500 × *g* for 5 min, washed 1X with PBS, and then pellets were snap frozen and stored at –80°C. Protein extraction, protein digestion and offline peptide fractionation was performed based on a protocol adapted from prior work(103). Briefly, cells were lysed in 8M urea buffer, reduced, alkylated, and digested using LysC and Trypsin proteases. The peptides were labeled with TMTpro 16 plex isobaric label reagent (Thermo Fisher Scientific) according to manufacturer’s protocol. The high-pH offline fractionation was used to generate 24 peptide pools. The deep-fractionated peptide samples were separated on an online nanoflow Easy-nLC-1200 system (Thermo Fisher Scientific) and analyzed on Orbitrap Exploris 480 mass spectrometer (Thermo Fisher Scientific). Each fraction (250 ng) was loaded on a pre-column (2 cm × 100µm I.D.) and separated on in-line 20 cm × 75 µm I.D. column (Reprosil-Pur Basic C18aq, Dr. Maisch GmbH, Germany) equilibrated in 0.1% formic acid (FA). Peptide separation was done at a flow rate of 200 nL/min over 110 min gradient time with different concentration of 90% acetonitrile solvent B (2-30% 87 min, 30-60% 6 min, 60-90% 7min and finally hold at 50% 10min). The heated column was maintained at 60°C. The mass spectrometer was operated in a data dependent mode with 2 s cycle time. The MS1 was done in Orbitrap (120000 resolution, scan range 375-1500 m/z, 50 ms injection time) followed by MS2 in Orbitrap at 30000 resolution (HCD 38%) with TurboTMT algorithm. Dynamic exclusion was set to 20 s and the isolation width was set to 0.7 m/z. The MS raw data processing, peptide validation, quantification and differential analysis was conducted as described before(104). The reverse decoys and common contaminants were added to the NCBI refseq human protein database (downloaded 2020.03.09) using Philosopher(105). Batch correction between multiplexes was performed using ComBat(106) as implemented in the R package Surrogate Variable Analysis (sva) version 3.44.0(107). Group differences were calculated using the moderated t-test as implemented in the R package limma(108) using default parameters with exception of robust = True, trend = True). Multiple-hypothesis testing correction was performed with the Benjamini–Hochberg procedure(109). Gene Set Enrichment Analysis (GSEA)(110) was performed using WebGestalt 2019(111) using the signed log *P* values from limma. Additional data analysis was performed using R version 4.2 and Python version 3.10(112), along with third-party scientific computing libraries NumPy(113) and Pandas(114).

### Human neutrophil flow cytometry

Peripheral human neutrophils were isolated as above and pretreated with 50 μg/mL of purified human THP or mock-treated, and incubated at 37°C in 5% CO2 for 30 min, and then incubated for an additional 2.5 h with PMA (25 nM). Cells were pelleted by centrifugation at 500 × *g* for 5 min, washed 1X with PBS, and resuspended in 50 μL of FACS buffer (1mM EDTA, 1% FBS, 0.1% sodium azide in PBS). The following antibodies (0.5 μg/mL) and dyes (concentrations provided below) were added: Anti-CD11b-FITC (clone M1/70, catalog no. 553310; BD Biosciences), anti-Ly6G-APC (clone 1A8, catalog no. 127614; BioLegend), Fc Block (catalog no. 564219; BD Pharmingen), Live/Dead Near IR (1:200 of stock, catalog no. L34975; Thermo Fisher Scientific), Sytox orange (100 nM, catalog no. S34861; Thermo Fisher Scientific), and Hoechst 33342 (200 nM, catalog no. 62249; Thermo Fisher Scientific). After a 30-minute incubation on ice, samples were washed 1X, resuspended in fresh FACS buffer, and run on a BD FACSCanto II (BD Biosciences). Samples were gated on unstained cells as described in **Fig. 4A** and positive signals were determined using single-stain controls, and data were analyzed with FlowJo version 10.9.0 (FlowJo LLC).

### Imaging flow cytometry and analyses

Peripheral human neutrophils were isolated as above and pretreated with 50 μg/mL of purified human THP, sialic acid (500 ng/mL, catalog no. A0812, Sigma Aldrich), or mock-treated, and incubated at 37°C in 5% CO2 for 30 min, and then incubated for an additional 2.5 h with PMA (25 nM). The following antibodies (0.5 μg/mL) and dyes (concentrations provided below) were added: Anti-MPO-FITC (clone M1/70, catalog no. 553310; BD Biosciences), Fc Block, Live/Dead Near IR (1:200 of stock, catalog no. L34975; Thermo Fisher Scientific), Sytox orange (100 nM, catalog no. S34861; Thermo Fisher Scientific), and Hoechst 33342 (200 nM, catalog no. 62249; Thermo Fisher Scientific). After a 30-minute incubation on ice, samples were washed 1X, resuspended in a 1:1 mixture of fresh FACS buffer and PBS. Cell morphology via imaging flow cytometry was assessed as described previously(76). An Amnis ImageStream X Mark II imaging flow cytometer was used for data acquisition with a 60X objective, low flow rate and high sensitivity, and 405, 488, 561 and 635 nm lasers set to 120, 150, 100 and 150LmW respectively. Data were analyzed using the IDEAS version 6.3 software package. Clipped images were retained due to the large size of NETs, and single stained controls were used for compensation. Masking was performed using the default “object (tight)” and “morphology” algorithms as described(76). Statistics reports were generated using IDEAS and processed IDEAS data analysis files (.daf) were then analyzed in FCS Express 7 to generate analysis plots.

### Statistics

Data were collected from at least two independent experiments unless indicated otherwise. Mean values from independent experiment replicates, or biological replicates, are represented by medians with interquartile ranges, or box-and-whisker plots with Tukey’s whiskers as indicated in figure legends. Experimental samples size (*n*) are indicated in figure legends. All data sets were subjected to D’Agostino & Pearson normality test to determine whether values displayed Gaussian distribution before selecting the appropriate parametric or non-parametric analyses. In the instances where *in vitro*, *ex vivo*, and *in vivo* experimental *n* were too small to determine normality, data were assumed non-parametric. For statistical comparisons of histopathology and urine sediment scores, mice were grouped into low (0–2) or high (>2) categories and frequencies were compared by Fisher’s exact test between WT and THP KO genotypes in both UPEC-infected and mock-infected conditions or IgG-treated and Ly6G-treated conditions respectively. UPEC urine and tissue burdens and THP urine levels were analyzed using two-tailed Mann-Whitney test. Immune cell populations, UPEC burdens in neutrophil depletion experiments, and flow cytometry of human and mouse neutrophil populations were analyzed using two-way ANOVA with Sidak’s multiple comparisons test or uncorrected Fisher’s Least Significant Difference (LSD) test as indicated in figure legends.

Imaging flow cytometry populations and murine sialic acid levels were compared using one-way ANOVA with Holm-Sidak’s multiple comparisons test. Proteomics data were analyzed by moderated t-test followed by multiple-hypothesis testing correction using the Benjamini– Hochberg procedure with a false discovery rate adjusted *P* <0.05. Statistical analyses were performed using GraphPad Prism, version 10.0.2 (GraphPad Software Inc., La Jolla, CA, USA). *P* values <0.05 were considered statistically significant.

### Study approval

Human peripheral blood and urine specimens for THP purification were obtained from healthy adult volunteers under approval from UC San Diego IRB (131002) and Baylor College of Medicine IRB (H-47537). All animal protocols and procedures were approved by UCSD and BCM Institutional Animal Care and Use Committees under protocols S00227M and AN-8233 respectively.

### Data availability

The mass spectrometry proteomics data have been deposited to the ProteomeXchange Consortium via the PRIDE partner repository(115) with the dataset identifier PXD045468. The ImageStream data is deposited in Figshare under project “ISX data files for THP NETosis Manuscript,” doi (https://doi.org/10.6084/m9.figshare.25013786, https://doi.org/10.6084/m9.figshare.25013774, https://doi.org/10.6084/m9.figshare.25013744, and https://doi.org/10.6084/m9.figshare.25013651).

## AUTHOR CONTRIBUTIONS

Research studies were designed by VN and KAP, VME, CC, CS, MEM, JJZ, and KAP conducted experiments and acquired data, data was analyzed by CC, AS, IC, and KAP, VME and KAP drafted the manuscript, and all authors contributed to manuscript review and edits.

## ACKNOWLEDGEMENTS

We are grateful to the vivarium staff at UCSD and BCM for animal husbandry and Dr. Jacqueline Kimmey for helpful discussions. Glycan analyses were performed at the UCSD GlycoAnalytics Core with assistance from Sulabah Argade, Mousumi Paulchakrabarti, and Biswa Choudhury. This project was supported by the BCM Mass Spectrometry Proteomics Core with assistance from Anna Malovannaya, Antrix Jain, and Mei Leng. BCM Mass Spectrometry Proteomics Core is supported by the Dan L. Duncan Comprehensive Cancer Center NIH award (P30 CA125123), CPRIT Core Facility Award (RP210227) and NIH High End Instrument award (S10 OD026804). This project was supported by the BCM Cytometry and Cell Sorting Core with funding from the CPRIT Core Facility Support Award (CPRIT-RP180672), the NIH (CA125123 and RR024574) and the assistance of Joel M. Sederstrom. VME, MEM, and JJZ were supported by NIH F31 awards (AI167547, AI167538, DK136201) respectively. Studies were supported NIH R01 (DK128053) and American Urological Association Research Scholar awards to KAP. The funders had no role in study design, data collection and interpretation, or the decision to submit the work for publication.

## Notes

### Competing Interest Statement

The authors have declared no competing interest.

## REFERENCES

1. Terlizzi ME, Gribaudo G, and Maffei ME. UroPathogenic Escherichia coli (UPEC) Infections: Virulence Factors, Bladder Responses, Antibiotic, and Non-antibiotic Antimicrobial Strategies. Front Microbiol. 2017;8:1566.

2. Tandogdu Z, and Wagenlehner FM. Global epidemiology of urinary tract infections. Curr Opin Infect Dis. 2016;29(1):73–9.

3. Yang X, Chen H, Zheng Y, Qu S, Wang H, and Yi F. Disease burden and long-term trends of urinary tract infections: A worldwide report. Front Public Health. 2022;10:888205.

4. Foxman B. Urinary tract infection syndromes: occurrence, recurrence, bacteriology, risk factors, and disease burden. Infect Dis Clin North Am. 2014;28(1):1–13.

5. Medina M, and Castillo-Pino E. An introduction to the epidemiology and burden of urinary tract infections. Ther Adv Urol. 2019;11:1756287219832172.

6. Ambite I, Nagy K, Godaly G, Puthia M, Wullt B, and Svanborg C. Susceptibility to Urinary Tract Infection: Benefits and Hazards of the Antibacterial Host Response. Microbiol Spectr. 2016;4(3).

7. Godaly G, Ambite I, and Svanborg C. Innate immunity and genetic determinants of urinary tract infection susceptibility. Curr Opin Infect Dis. 2015;28(1):88–96.

8. Jaillon S, Moalli F, Ragnarsdottir B, Bonavita E, Puthia M, Riva F, et al. The humoral pattern recognition molecule PTX3 is a key component of innate immunity against urinary tract infection. Immunity. 2014;40(4):621–32.

9. Chu CM, and Lowder JL. Diagnosis and treatment of urinary tract infections across age groups. Am J Obstet Gynecol. 2018;219(1):40–51.

10. Bergsten G, Samuelsson M, Wullt B, Leijonhufvud I, Fischer H, and Svanborg C. PapG-dependent adherence breaks mucosal inertia and triggers the innate host response. J Infect Dis. 2004;189(9):1734–42.

11. Sundac L, Dando SJ, Sullivan MJ, Derrington P, Gerrard J, and Ulett GC. Protein-based profiling of the immune response to uropathogenic Escherichia coli in adult patients immediately following hospital admission for acute cystitis. Pathog Dis. 2016;74(6).

12. Isaacson B, Hadad T, Glasner A, Gur C, Granot Z, Bachrach G, et al. Stromal Cell-Derived Factor 1 Mediates Immune Cell Attraction upon Urinary Tract Infection. Cell Rep. 2017;20(1):40–7.

13. Ingersoll MA, Kline KA, Nielsen HV, and Hultgren SJ. G-CSF induction early in uropathogenic Escherichia coli infection of the urinary tract modulates host immunity. Cell Microbiol. 2008;10(12):2568–78.

14. Haraoka M, Hang L, Frendeus B, Godaly G, Burdick M, Strieter R, et al. Neutrophil recruitment and resistance to urinary tract infection. J Infect Dis. 1999;180(4):1220–9.

15. Frendeus B, Godaly G, Hang L, Karpman D, Lundstedt AC, and Svanborg C. Interleukin 8 receptor deficiency confers susceptibility to acute experimental pyelonephritis and may have a human counterpart. J Exp Med. 2000;192(6):881–90.

16. Svensson M, Irjala H, Alm P, Holmqvist B, Lundstedt AC, and Svanborg C. Natural history of renal scarring in susceptible mIL-8Rh-/-mice. Kidney Int. 2005;67(1):103–10.

17. Hannan TJ, Roberts PL, Riehl TE, van der Post S, Binkley JM, Schwartz DJ, et al. Inhibition of Cyclooxygenase-2 Prevents Chronic and Recurrent Cystitis. EBioMedicine. 2014;1(1):46–57.

18. Chromek M, Slamova Z, Bergman P, Kovacs L, Podracka L, Ehren I, et al. The antimicrobial peptide cathelicidin protects the urinary tract against invasive bacterial infection. Nat Med. 2006;12(6):636–41.

19. Patras KA, Coady A, Babu P, Shing SR, Ha AD, Rooholfada E, et al. Host Cathelicidin Exacerbates Group B Streptococcus Urinary Tract Infection. mSphere. 2020;5(2).

20. Patras KA, Ha AD, Rooholfada E, Olson J, Ramachandra Rao SP, Lin AE, et al. Augmentation of Urinary Lactoferrin Enhances Host Innate Immune Clearance of Uropathogenic Escherichia coli. J Innate Immun. 2019;11(6):481–95.

21. Arao S, Matsuura S, Nonomura M, Miki K, Kabasawa K, and Nakanishi H. Measurement of urinary lactoferrin as a marker of urinary tract infection. J Clin Microbiol. 1999;37(3):553–7.

22. Schiwon M, Weisheit C, Franken L, Gutweiler S, Dixit A, Meyer-Schwesinger C, et al. Crosstalk between sentinel and helper macrophages permits neutrophil migration into infected uroepithelium. Cell. 2014;156(3):456–68.

23. Mundi H, Bjorksten B, Svanborg C, Ohman L, and Dahlgren C. Extracellular release of reactive oxygen species from human neutrophils upon interaction with Escherichia coli strains causing renal scarring. Infect Immun. 1991;59(11):4168–72.

24. Gupta A, Sharma S, Nain CK, Sharma BK, and Ganguly NK. Reactive oxygen species-mediated tissue injury in experimental ascending pyelonephritis. Kidney Int. 1996;49(1):26–33.

25. Condron C, Toomey D, Casey RG, Shaffii M, Creagh T, and Bouchier-Hayes D. Neutrophil bactericidal function is defective in patients with recurrent urinary tract infections. Urol Res. 2003;31(5):329–34.

26. Brinkmann V, Reichard U, Goosmann C, Fauler B, Uhlemann Y, Weiss DS, et al. Neutrophil extracellular traps kill bacteria. Science. 2004;303(5663):1532–5.

27. Metzler KD, Fuchs TA, Nauseef WM, Reumaux D, Roesler J, Schulze I, et al. Myeloperoxidase is required for neutrophil extracellular trap formation: implications for innate immunity. Blood. 2011;117(3):953–9.

28. Hoppenbrouwers T, Autar ASA, Sultan AR, Abraham TE, van Cappellen WA, Houtsmuller AB, et al. In vitro induction of NETosis: Comprehensive live imaging comparison and systematic review. PLoS One. 2017;12(5):e0176472.

29. Papayannopoulos V. Neutrophil extracellular traps in immunity and disease. Nat Rev Immunol. 2018;18(2):134–47.

30. Tan C, Aziz M, and Wang P. The vitals of NETs. J Leukoc Biol. 2021;110(4):797–808.

31. Yousefi S, Mihalache C, Kozlowski E, Schmid I, and Simon HU. Viable neutrophils release mitochondrial DNA to form neutrophil extracellular traps. Cell Death Differ. 2009;16(11):1438–44.

32. Yipp BG, Petri B, Salina D, Jenne CN, Scott BN, Zbytnuik LD, et al. Infection-induced NETosis is a dynamic process involving neutrophil multitasking in vivo. Nat Med. 2012;18(9):1386–93.

33. Pilsczek FH, Salina D, Poon KK, Fahey C, Yipp BG, Sibley CD, et al. A novel mechanism of rapid nuclear neutrophil extracellular trap formation in response to Staphylococcus aureus. J Immunol. 2010;185(12):7413–25.

34. Yu Y, Kwon K, Tsitrin T, Bekele S, Sikorski P, Nelson KE, et al. Characterization of Early-Phase Neutrophil Extracellular Traps in Urinary Tract Infections. PLoS Pathog. 2017;13(1):e1006151.

35. Yu Y, Kwon K, and Pieper R. Detection of Neutrophil Extracellular Traps in Urine. Methods Mol Biol. 2019;2021:241–57.

36. Sharma K, Dhar N, Thacker VV, Simonet TM, Signorino-Gelo F, Knott GW, et al. Dynamic persistence of UPEC intracellular bacterial communities in a human bladder-chip model of urinary tract infection. Elife. 2021;10.

37. Mo L, Huang HY, Zhu XH, Shapiro E, Hasty DL, and Wu XR. Tamm-Horsfall protein is a critical renal defense factor protecting against calcium oxalate crystal formation. Kidney Int. 2004;66(3):1159–66.

38. Liu Y, Goldfarb DS, El-Achkar TM, Lieske JC, and Wu XR. Tamm-Horsfall protein/uromodulin deficiency elicits tubular compensatory responses leading to hypertension and hyperuricemia. Am J Physiol Renal Physiol. 2018;314(6):F1062–F76.

39. Bates JM, Raffi HM, Prasadan K, Mascarenhas R, Laszik Z, Maeda N, et al. Tamm-Horsfall protein knockout mice are more prone to urinary tract infection: rapid communication. Kidney Int. 2004;65(3):791–7.

40. Mo L, Zhu XH, Huang HY, Shapiro E, Hasty DL, and Wu XR. Ablation of the Tamm-Horsfall protein gene increases susceptibility of mice to bladder colonization by type 1-fimbriated Escherichia coli. Am J Physiol Renal Physiol. 2004;286(4):F795–802.

41. Raffi HS, Bates JM, Jr., Laszik Z, and Kumar S. Tamm-horsfall protein protects against urinary tract infection by proteus mirabilis. J Urol. 2009;181(5):2332–8.

42. Coady A, Ramos AR, Olson J, Nizet V, and Patras KA. Tamm-Horsfall Protein Protects the Urinary Tract against Candida albicans. Infect Immun. 2018;86(12).

43. Pak J, Pu Y, Zhang ZT, Hasty DL, and Wu XR. Tamm-Horsfall protein binds to type 1 fimbriated Escherichia coli and prevents E. coli from binding to uroplakin Ia and Ib receptors. J Biol Chem. 2001;276(13):9924–30.

44. Leeker A, Kreft B, Sandmann J, Bates J, Wasenauer G, Muller H, et al. Tamm-Horsfall protein inhibits binding of S-and P-fimbriated Escherichia coli to human renal tubular epithelial cells. Exp Nephrol. 1997;5(1):38–46.

45. Harjai K, Mittal R, Chhibber S, and Sharma S. Contribution of Tamm-Horsfall protein to virulence of Pseudomonas aeruginosa in urinary tract infection. Microbes Infect. 2005;7(1):132–7.

46. Weiss GL, Stanisich JJ, Sauer MM, Lin CW, Eras J, Zyla DS, et al. Architecture and function of human uromodulin filaments in urinary tract infections. Science. 2020;369(6506):1005-10.

47. Yu CL, Lin WM, Liao TS, Tsai CY, Sun KH, and Chen KH. Tamm-Horsfall glycoprotein (THG) purified from normal human pregnancy urine increases phagocytosis, complement receptor expressions and arachidonic acid metabolism of polymorphonuclear neutrophils. Immunopharmacology. 1992;24(3):181–90.

48. Su SJ, Chang KL, Lin TM, Huang YH, and Yeh TM. Uromodulin and Tamm-Horsfall protein induce human monocytes to secrete TNF and express tissue factor. J Immunol. 1997;158(7):3449–56.

49. Su SJ, and Yeh TM. The dynamic responses of pro-inflammatory and anti-inflammatory cytokines of human mononuclear cells induced by uromodulin. Life Sci. 1999;65(24):2581–90.

50. Patras KA, Coady A, Olson J, Ali SR, RamachandraRao SP, Kumar S, et al. Tamm-Horsfall glycoprotein engages human Siglec-9 to modulate neutrophil activation in the urinary tract. Immunol Cell Biol. 2017;95(10):960–5.

51. Liu Y, El-Achkar TM, and Wu XR. Tamm-Horsfall protein regulates circulating and renal cytokines by affecting glomerular filtration rate and acting as a urinary cytokine trap. J Biol Chem. 2012;287(20):16365–78.

52. El-Achkar TM, Wu XR, Rauchman M, McCracken R, Kiefer S, and Dagher PC. Tamm-Horsfall protein protects the kidney from ischemic injury by decreasing inflammation and altering TLR4 expression. Am J Physiol Renal Physiol. 2008;295(2):F534–44.

53. Micanovic R, Chitteti BR, Dagher PC, Srour EF, Khan S, Hato T, et al. Tamm-Horsfall Protein Regulates Granulopoiesis and Systemic Neutrophil Homeostasis. J Am Soc Nephrol. 2015;26(9):2172–82.

54. Zulk JJ, Clark JR, Ottinger S, Ballard MB, Mejia ME, Mercado-Evans V, et al. Phage Resistance Accompanies Reduced Fitness of Uropathogenic Escherichia coli in the Urinary Environment. mSphere. 2022;7(4):e0034522.

55. Dou W, Thompson-Jaeger S, Laulederkind SJ, Becker JW, Montgomery J, Ruiz-Bustos E, et al. Defective expression of Tamm-Horsfall protein/uromodulin in COX-2-deficient mice increases their susceptibility to urinary tract infections. Am J Physiol Renal Physiol. 2005;289(1):F49–60.

56. Mayorek N, Naftali-Shani N, and Grunewald M. Diclofenac inhibits tumor growth in a murine model of pancreatic cancer by modulation of VEGF levels and arginase activity. PLoS One. 2010;5(9):e12715.

57. Ghirotto S, Tassi F, Barbujani G, Pattini L, Hayward C, Vollenweider P, et al. The Uromodulin Gene Locus Shows Evidence of Pathogen Adaptation through Human Evolution. J Am Soc Nephrol. 2016;27(10):2983–96.

58. Garimella PS, Bartz TM, Ix JH, Chonchol M, Shlipak MG, Devarajan P, et al. Urinary Uromodulin and Risk of Urinary Tract Infections: The Cardiovascular Health Study. Am J Kidney Dis. 2017;69(6):744–51.

59. Stahl K, Beneke J, Haller H, Gwinner W, and Schiffer M. Reduced Urinary Uromodulin (UMOD)-Levels Are associated With Urinary Tract Infections (UTI) After Renal Transplantion. Am J Transplant. 2015(15):suppl 3.

60. Reinhart HH, Spencer JR, Zaki NF, and Sobel JD. Quantitation of urinary Tamm-Horsfall protein in children with urinary tract infection. Eur Urol. 1992;22(3):194–9.

61. Parsons CL, Proctor J, Teichman JS, Nickel JC, Davis E, Evans R, et al. A multi-site study confirms abnormal glycosylation in the Tamm-Horsfall protein of patients with interstitial cystitis. J Urol. 2011;186(1):112–6.

62. Argade S, Chen T, Shaw T, Berecz Z, Shi W, Choudhury B, et al. An evaluation of Tamm-Horsfall protein glycans in kidney stone formers using novel techniques. Urolithiasis. 2015;43(4):303–12.

63. Li H, Kostel SA, DiMartino SE, Hashemi Gheinani A, Froehlich JW, and Lee RS. Uromodulin Isolation and Its N-Glycosylation Analysis by NanoLC-MS/MS. J Proteome Res. 2021;20(5):2662–72.

64. Singhal A, Yadav S, Chandra T, Mulay SR, Gaikwad AN, and Kumar S. An Imaging and Computational Algorithm for Efficient Identification and Quantification of Neutrophil Extracellular Traps. Cells. 2022;11(2).

65. Zharkova O, Tay SH, Lee HY, Shubhita T, Ong WY, Lateef A, et al. A Flow Cytometry-Based Assay for High-Throughput Detection and Quantification of Neutrophil Extracellular Traps in Mixed Cell Populations. Cytometry A. 2019;95(3):268–78.

66. Masuda S, Shimizu S, Matsuo J, Nishibata Y, Kusunoki Y, Hattanda F, et al. Measurement of NET formation in vitro and in vivo by flow cytometry. Cytometry A. 2017;91(8):822–9.

67. Perfetto SP, Chattopadhyay PK, Lamoreaux L, Nguyen R, Ambrozak D, Koup RA, et al. Amine-reactive dyes for dead cell discrimination in fixed samples. Curr Protoc Cytom. 2010;Chapter 9:Unit 9 34.

68. Yousefi S, Simon D, Stojkov D, Karsonova A, Karaulov A, and Simon HU. In vivo evidence for extracellular DNA trap formation. Cell Death Dis. 2020;11(4):300.

69. Vorobjeva NV, and Chernyak BV. NETosis: Molecular Mechanisms, Role in Physiology and Pathology. Biochemistry (Mosc). 2020;85(10):1178–90.

70. Swensen AC, He J, Fang AC, Ye Y, Nicora CD, Shi T, et al. A Comprehensive Urine Proteome Database Generated From Patients With Various Renal Conditions and Prostate Cancer. Front Med (Lausanne). 2021;8:548212.

71. Parker H, Dragunow M, Hampton MB, Kettle AJ, and Winterbourn CC. Requirements for NADPH oxidase and myeloperoxidase in neutrophil extracellular trap formation differ depending on the stimulus. J Leukoc Biol. 2012;92(4):841–9.

72. Fuchs TA, Abed U, Goosmann C, Hurwitz R, Schulze I, Wahn V, et al. Novel cell death program leads to neutrophil extracellular traps. J Cell Biol. 2007;176(2):231–41.

73. Thiam HR, Wong SL, Qiu R, Kittisopikul M, Vahabikashi A, Goldman AE, et al. NETosis proceeds by cytoskeleton and endomembrane disassembly and PAD4-mediated chromatin decondensation and nuclear envelope rupture. Proc Natl Acad Sci U S A. 2020;117(13):7326–37.

74. Remijsen Q, Vanden Berghe T, Wirawan E, Asselbergh B, Parthoens E, De Rycke R, et al. Neutrophil extracellular trap cell death requires both autophagy and superoxide generation. Cell Res. 2011;21(2):290–304.

75. Neubert E, Meyer D, Rocca F, Gunay G, Kwaczala-Tessmann A, Grandke J, et al. Chromatin swelling drives neutrophil extracellular trap release. Nat Commun. 2018;9(1):3767.

76. Lelliott PM, Momota M, Lee MSJ, Kuroda E, Iijima N, Ishii KJ, et al. Rapid Quantification of NETs In Vitro and in Whole Blood Samples by Imaging Flow Cytometry. Cytometry A. 2019;95(5):565–78.

77. Barbu EA, Dominical VM, Mendelsohn L, and Thein SL. Detection and Quantification of Histone H4 Citrullination in Early NETosis With Image Flow Cytometry Version 4. Front Immunol. 2020;11:1335.

78. Raffi HS, Bates JM, Jr., Laszik Z, and Kumar S. Tamm-Horsfall protein acts as a general host-defense factor against bacterial cystitis. Am J Nephrol. 2005;25(6):570–8.

79. El-Achkar TM, McCracken R, Rauchman M, Heitmeier MR, Al-Aly Z, Dagher PC, et al. Tamm-Horsfall protein-deficient thick ascending limbs promote injury to neighboring S3 segments in an MIP-2-dependent mechanism. Am J Physiol Renal Physiol. 2011;300(4):F999–1007.

80. Liu Y, Mo L, Goldfarb DS, Evan AP, Liang F, Khan SR, et al. Progressive renal papillary calcification and ureteral stone formation in mice deficient for Tamm-Horsfall protein. Am J Physiol Renal Physiol. 2010;299(3):F469–78.

81. Olczak T, Olczak M, Kubicz A, Dulawa J, and Kokot F. Composition of the sugar moiety of Tamm-Horsfall protein in patients with urinary diseases. Int J Clin Lab Res. 1999;29(2):68–74.

82. Prasadan K, Bates J, Badgett A, Dell M, Sukhatme V, Yu H, et al. Nucleotide sequence and peptide motifs of mouse uromodulin (Tamm-Horsfall protein)--the most abundant protein in mammalian urine. Biochim Biophys Acta. 1995;1260(3):328–32.

83. Hedlund M, Tangvoranuntakul P, Takematsu H, Long JM, Housley GD, Kozutsumi Y, et al. N-glycolylneuraminic acid deficiency in mice: implications for human biology and evolution. Mol Cell Biol. 2007;27(12):4340–6.

84. Fischer H, Lutay N, Ragnarsdottir B, Yadav M, Jonsson K, Urbano A, et al. Pathogen specific, IRF3-dependent signaling and innate resistance to human kidney infection. PLoS Pathog. 2010;6(9):e1001109.

85. Moreira-Teixeira L, Stimpson PJ, Stavropoulos E, Hadebe S, Chakravarty P, Ioannou M, et al. Type I IFN exacerbates disease in tuberculosis-susceptible mice by inducing neutrophil-mediated lung inflammation and NETosis. Nat Commun. 2020;11(1):5566.

86. Pylaeva E, Bordbari S, Spyra I, Decker AS, Haussler S, Vybornov V, et al. Detrimental Effect of Type I IFNs During Acute Lung Infection With Pseudomonas aeruginosa Is Mediated Through the Stimulation of Neutrophil NETosis. Front Immunol. 2019;10:2190.

87. Krivosikova K, Supcikova N, Gaal Kovalcikova A, Janko J, Pastorek M, Celec P, et al. Neutrophil extracellular traps in urinary tract infection. Front Pediatr. 2023;11:1154139.

88. Petretto A, Bruschi M, Pratesi F, Croia C, Candiano G, Ghiggeri G, et al. Neutrophil extracellular traps (NET) induced by different stimuli: A comparative proteomic analysis. PLoS One. 2019;14(7):e0218946.

89. Wang X, Zhao J, Cai C, Tang X, Fu L, Zhang A, et al. A Label-Free Quantitative Proteomic Analysis of Mouse Neutrophil Extracellular Trap Formation Induced by Streptococcus suis or Phorbol Myristate Acetate (PMA). Front Immunol. 2018;9:2615.

90. Aquino EN, Neves AC, Santos KC, Uribe CE, Souza PE, Correa JR, et al. Proteomic Analysis of Neutrophil Priming by PAF. Protein Pept Lett. 2016;23(2):142–51.

91. Aarts CEM, Downes K, Hoogendijk AJ, Sprenkeler EGG, Gazendam RP, Favier R, et al. Neutrophil specific granule and NETosis defects in gray platelet syndrome. Blood Adv. 2021;5(2):549–64.

92. Khan MA, and Palaniyar N. Transcriptional firing helps to drive NETosis. Sci Rep. 2017;7:41749.

93. Bjornsdottir H, Welin A, Michaelsson E, Osla V, Berg S, Christenson K, et al. Neutrophil NET formation is regulated from the inside by myeloperoxidase-processed reactive oxygen species. Free Radic Biol Med. 2015;89:1024–35.

94. Stojkov D, Amini P, Oberson K, Sokollik C, Duppenthaler A, Simon HU, et al. ROS and glutathionylation balance cytoskeletal dynamics in neutrophil extracellular trap formation. J Cell Biol. 2017;216(12):4073–90.

95. Douda DN, Khan MA, Grasemann H, and Palaniyar N. SK3 channel and mitochondrial ROS mediate NADPH oxidase-independent NETosis induced by calcium influx. Proc Natl Acad Sci U S A. 2015;112(9):2817–22.

96. von Gunten S, Yousefi S, Seitz M, Jakob SM, Schaffner T, Seger R, et al. Siglec-9 transduces apoptotic and nonapoptotic death signals into neutrophils depending on the proinflammatory cytokine environment. Blood. 2005;106(4):1423–31.

97. Zhao W, Fogg DK, and Kaplan MJ. A novel image-based quantitative method for the characterization of NETosis. J Immunol Methods. 2015;423:104–10.

98. Secundino I, Lizcano A, Roupe KM, Wang X, Cole JN, Olson J, et al. Host and pathogen hyaluronan signal through human siglec-9 to suppress neutrophil activation. J Mol Med (Berl). 2016;94(2):219–33.

99. Khatua B, Bhattacharya K, and Mandal C. Sialoglycoproteins adsorbed by Pseudomonas aeruginosa facilitate their survival by impeding neutrophil extracellular trap through siglec-9. J Leukoc Biol. 2012;91(4):641–55.

100. Scharf B, Sendker J, Dobrindt U, and Hensel A. Influence of Cranberry Extract on Tamm-Horsfall Protein in Human Urine and its Antiadhesive Activity Against Uropathogenic Escherichia coli. Planta Med. 2019;85(2):126–38.

101. Mo B, Sendker J, Herrmann F, Nowak S, and Hensel A. Aqueous extract from Equisetum arvense stimulates the secretion of Tamm-Horsfall protein in human urine after oral intake. Phytomedicine. 2022;104:154302.

102. Mulvey MA, Schilling JD, and Hultgren SJ. Establishment of a persistent Escherichia coli reservoir during the acute phase of a bladder infection. Infect Immun. 2001;69(7):4572–9.

103. Mertins P, Yang F, Liu T, Mani DR, Petyuk VA, Gillette MA, et al. Ischemia in tumors induces early and sustained phosphorylation changes in stress kinase pathways but does not affect global protein levels. Mol Cell Proteomics. 2014;13(7):1690–704.

104. Nozawa K, Garcia TX, Kent K, Leng M, Jain A, Malovannaya A, et al. Testis-specific serine kinase 3 is required for sperm morphogenesis and male fertility. Andrology. 2023;11(5):826–39.

105. da Veiga Leprevost F, Haynes SE, Avtonomov DM, Chang HY, Shanmugam AK, Mellacheruvu D, et al. Philosopher: a versatile toolkit for shotgun proteomics data analysis. Nat Methods. 2020;17(9):869–70.

106. Johnson WE, Li C, and Rabinovic A. Adjusting batch effects in microarray expression data using empirical Bayes methods. Biostatistics. 2007;8(1):118–27.

107. Leek J, Johnson W, Parker H, Fertig E, Jaffe A, Zhang Y, et al. Surrogate Variable Analysis (sva) version 3.44.0. 2022.

108. Ritchie ME, Phipson B, Wu D, Hu Y, Law CW, Shi W, et al. limma powers differential expression analyses for RNA-sequencing and microarray studies. Nucleic Acids Res. 2015;43(7):e47.

109. Benjamini Y, and Hochberg Y. Controlling the False Discovery Rate: A Practical and Powerful Approach to Multiple Testing. J R Statist Soc B. 1995;57(1):289–300.

110. Subramanian A, Tamayo P, Mootha VK, Mukherjee S, Ebert BL, Gillette MA, et al. Gene set enrichment analysis: a knowledge-based approach for interpreting genome-wide expression profiles. Proc Natl Acad Sci U S A. 2005;102(43):15545–50.

111. Liao Y, Wang J, Jaehnig EJ, Shi Z, and Zhang B. WebGestalt 2019: gene set analysis toolkit with revamped UIs and APIs. Nucleic Acids Res. 2019;47(W1):W199–W205.

112. Anon. Python Package Index – PyPI. 2021.

113. Harris CR, Millman KJ, van der Walt SJ, Gommers R, Virtanen P, Cournapeau D, et al. Array programming with NumPy. Nature. 2020;585(7825):357–62.

114. team Tpd. pandas-dev/pandas: Pandas. 2020.

115. Perez-Riverol Y, Bai J, Bandla C, Garcia-Seisdedos D, Hewapathirana S, Kamatchinathan S, et al. The PRIDE database resources in 2022: a hub for mass spectrometry-based proteomics evidences. Nucleic Acids Res. 2022;50(D1):D543–D52.

