## Supplemental Material for "Tamm-Horsfall protein augments neutrophil NETosis during urinary tract infection"

**Contents:**

Supplementary Figures 1-6

Supplementary Tables 1-2

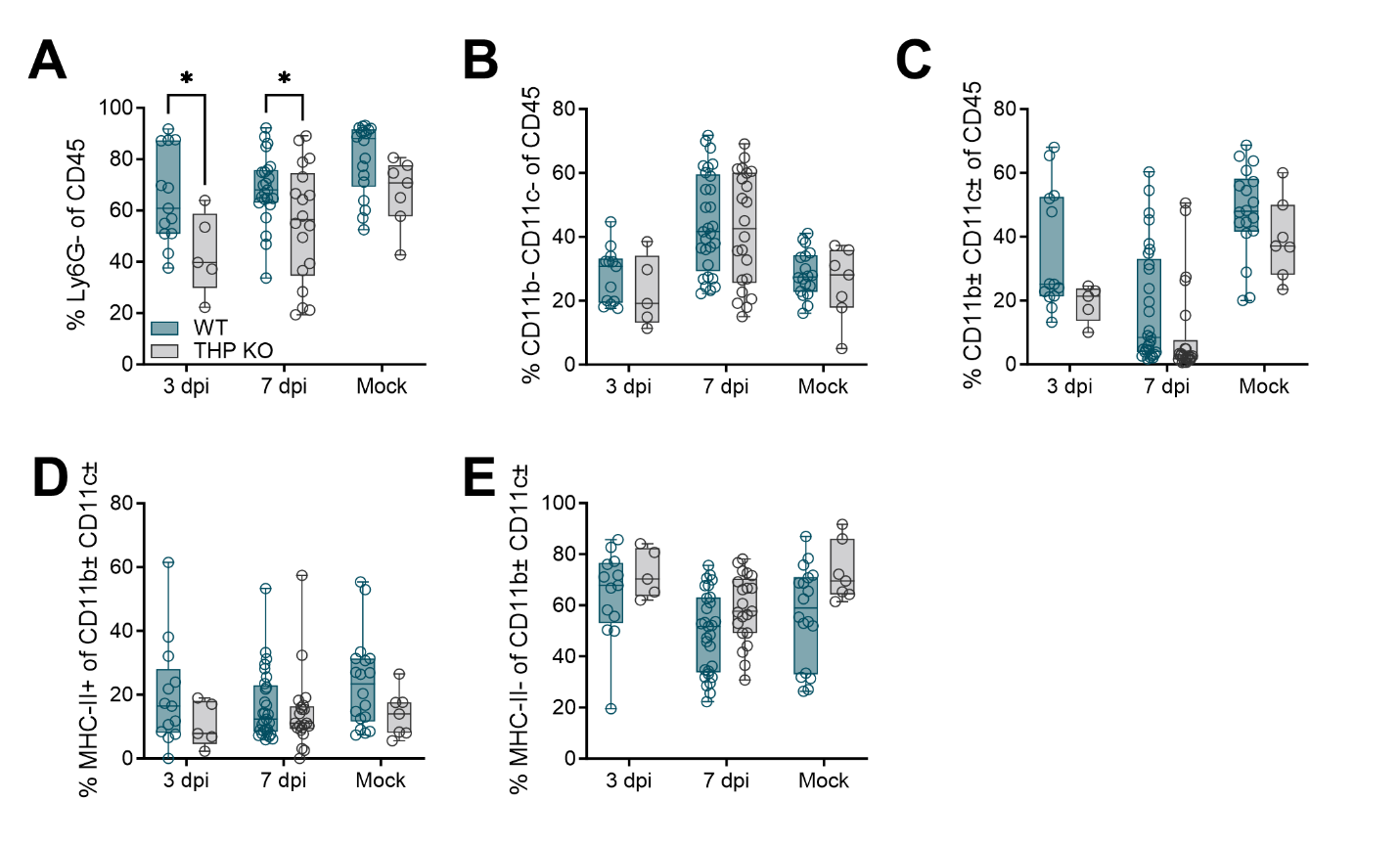
**Supplemental Figure 1. Impact of THP on other bladder immune populations during UTI.** Wild type (WT) and THP knockout (THP KO) mice were transurethrally infected with 10^8^ CFU of UPEC strain UTI89 or mock-infected as a control. Frequency of Ly6G- cells (A), CD11b-CD11c- (lymphocytes) cells (B), and CD11b^var^CD11c^var^ (myeloid) cells (C) as a percentage of total bladder CD45+ cells. Frequency of MHC-II+ (antigen presenting cells) (D) and MHC-II- (other myeloid) cells (E) as a percentage total myeloid (CD11b^var^CD11c^var^) cells. Experiments were performed at least two times with data combined, *n* = 5-23/group. Box and whisker plots extend from 25th to 75th percentiles and show all points. Data was analyzed by two-way ANOVA with Sidak’s multiple comparisons test. * *P* < 0.05. Supplemental data to **Figure 2**.

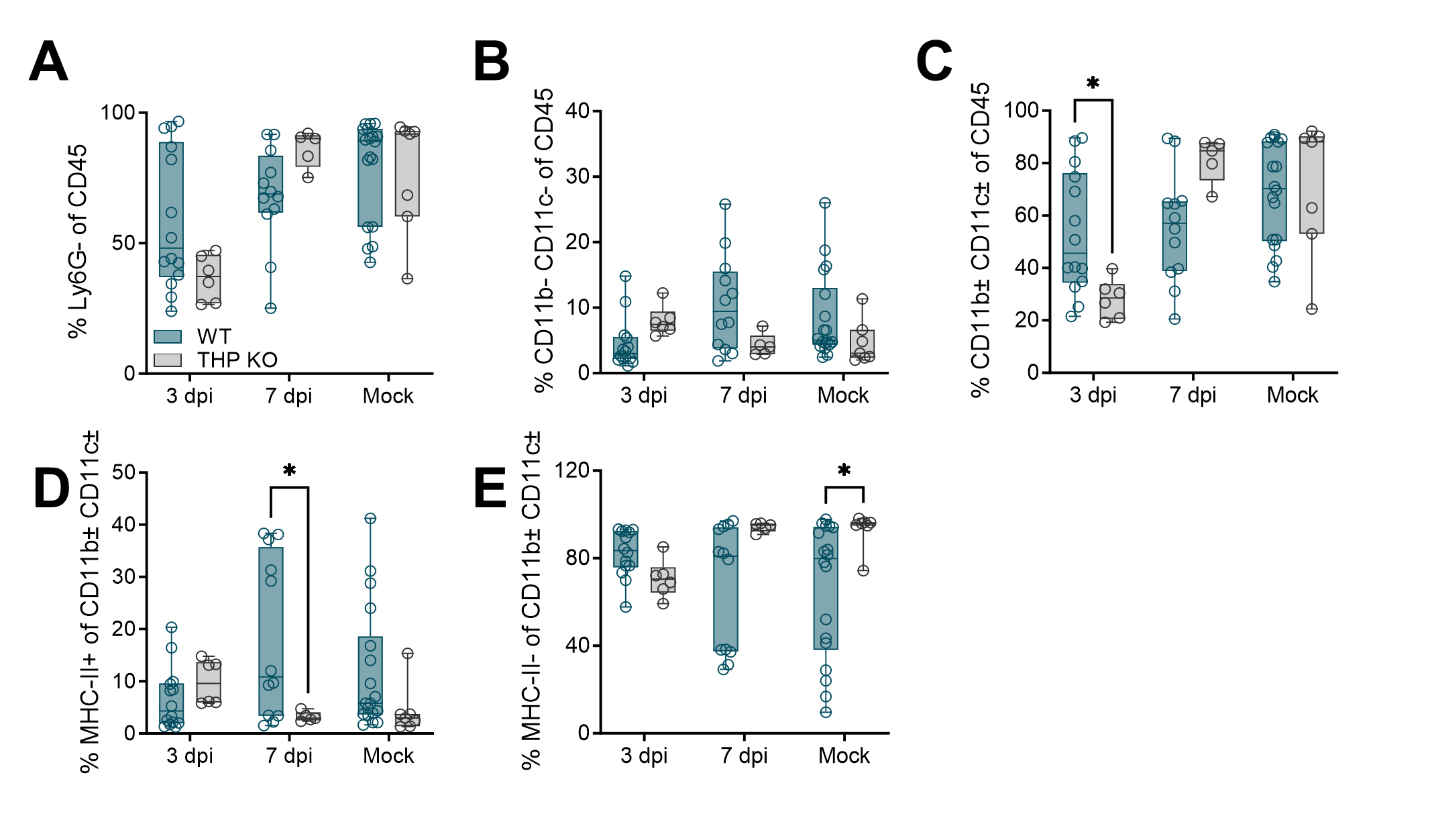

**Supplemental Figure 2. Impact of THP on other kidney immune populations during UTI.** Wild type (WT) and THP knockout (THP KO) mice were transurethrally infected with 10^8^ CFU of UPEC strain UTI89 or mock-infected as a control. Frequency of Ly6G- cells (A), CD11b-CD11c- (lymphocytes) cells (B), and CD11b^var^CD11c^var^ (myeloid) cells (C) as a percentage of total kidney CD45+ cells. Frequency of MHC-II+ (antigen presenting cells) (D) and MHC-II- (other myeloid) cells (E) as a percentage total myeloid (CD11b^var^CD11c^var^) cells. Experiments were performed at least two times with data combined, *n* = 5-23/group. Box and whisker plots extend from 25th to 75th percentiles and show all points. Data was analyzed by two-way ANOVA with Sidak’s multiple comparisons test. * *P* < 0.05. Supplemental data to **Figure 2**.

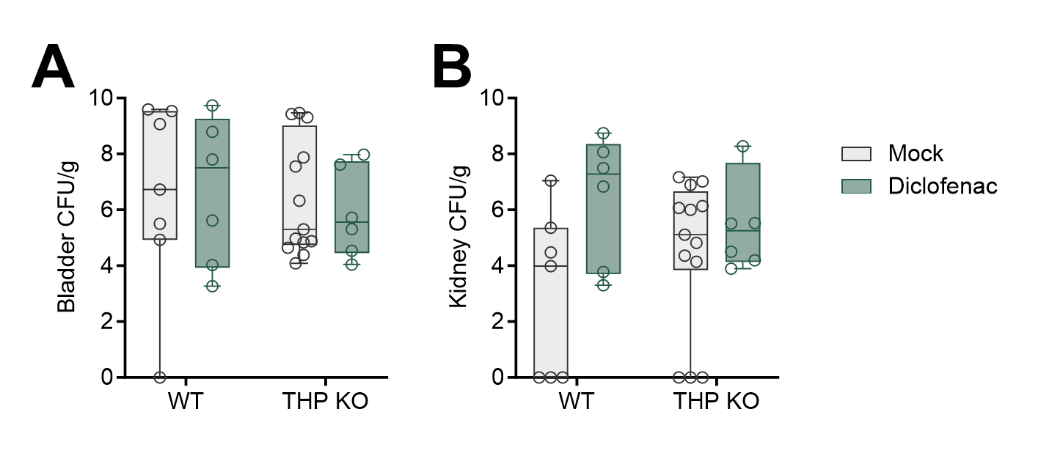
**Supplemental Figure 3. Cox-2 inhibitor Diclonfenac does not alter bacterial burdens in WT or THP KO mice**. Wild type (WT) and THP knockout (THP KO) mice were transurethrally infected with 10^8^ CFU of UPEC strain UTI89. To inhibit COX-2, mice were administered diclofenac in the drinking water beginning on day 0 and maintained through day 6 post-infection. Bladder (**A**) and kidney (**B**) UPEC burdens at 7 days post-infection. Experiments were performed at least two times with data combined, *n* = 6-13/group. Box and whisker plots extend from 25th to 75th percentiles and show all points. Data was analyzed by two-way ANOVA with Sidak’s multiple comparisons test and all comparisons were determine not significant (*P* > 0.05). Supplemental data to **Figure 2**.

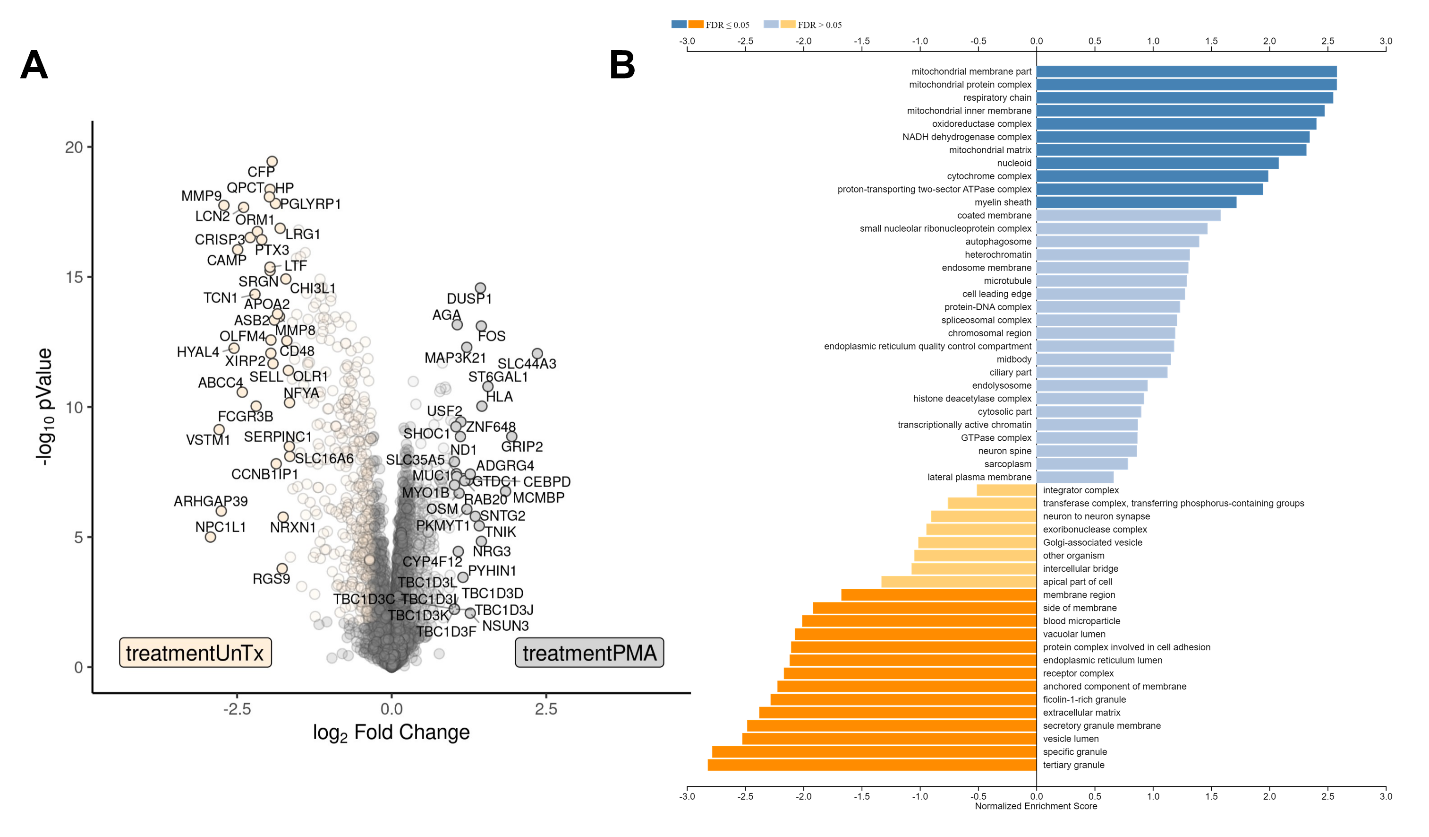

**Supplemental Figure 4. PMA stimulation alters the neutrophil proteome.** Peripheral human neutrophils were isolated, pretreated with 50 μg/mL THP, and stimulated phorbol 12-myristate 13-acetate (PMA) for 2.5 hours. Volcano plot (**A**) and gene set enrichment analysis (**E**) of differentially identified proteins in untreated vs. PMA stimulated samples. NES = Normalized Enrichment Score. Experiments were performed as part of one independent experiment, *n* = 4 donors. Differential proteins were identified via Log_2_ fold change >1.25 and moderated t-test followed by multiple-hypothesis testing correction using the Benjamini–Hochberg procedure with a false discovery rate adjusted *P* <0.05. GSEA was performed with a gene set minimum of 10, a gene set maximum of 500, 2,000 permutations using the gene ontology cellular component gene sets. Supplemental data to proteomic analyses in **Figure 5**.

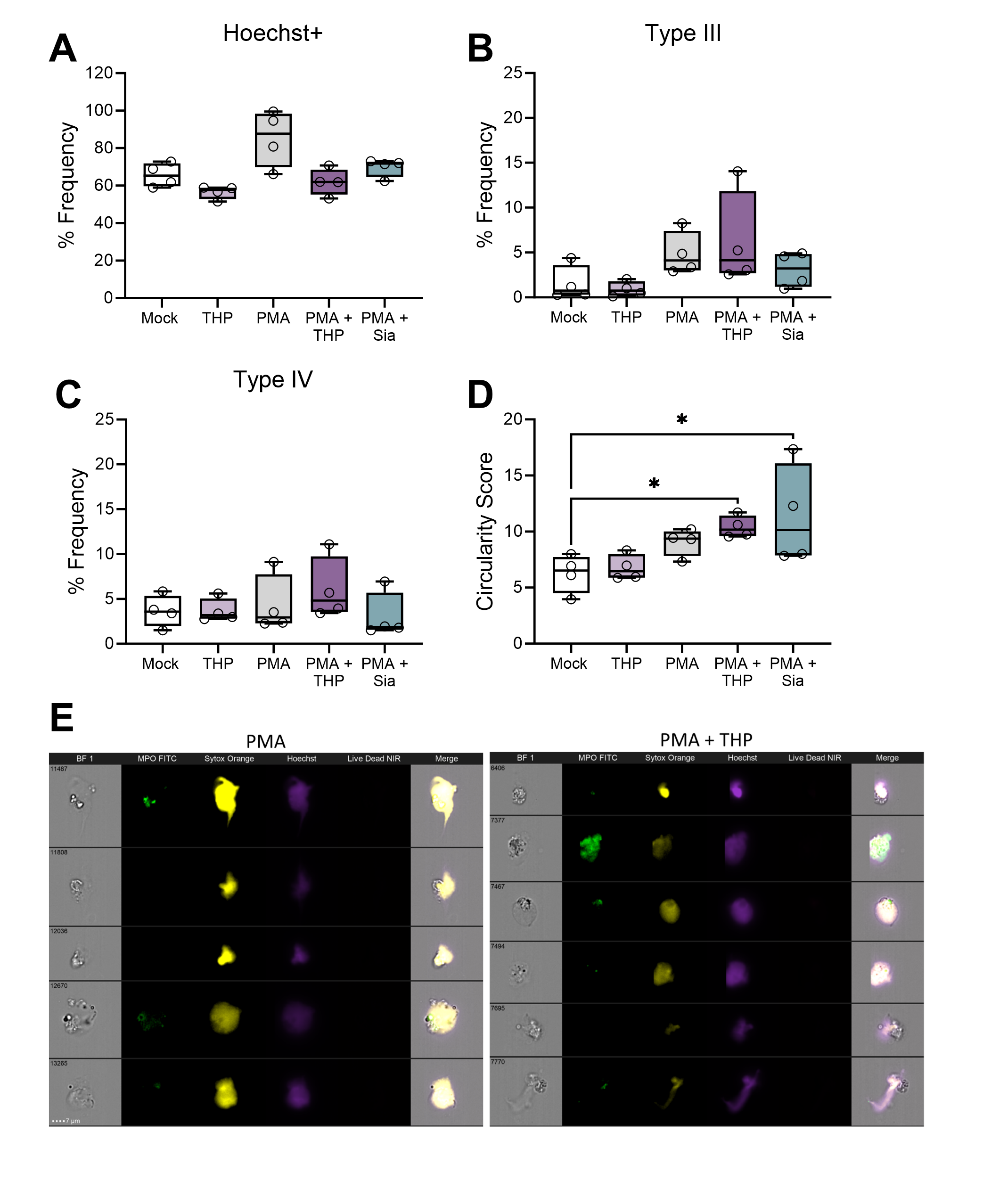
**Supplemental Figure 5. Additional effects of THP and sialic acid treatment on cellular morphologies as determined by imaging flow cytometry.** Peripheral human neutrophils were isolated, pretreated with THP or sialic acid, and stimulated phorbol 12-myristate 13-acetate (PMA) for 2.5 hours. Cells were stained with anti-MPO FITC, Sytox Orange (non-membrane permeable nucleic acid dye), Hoechst (membrane-permeable nucleic acid dye), and Live/Dead stain (non-membrane permeable amine-reactive dye) and visualized for fluorescence and brightfield (BF) images on an imaging flow cytometer. (**A**) Frequency of Hoechst+ events across treatment groups. (**B**) Frequency of NETs (Type III) gated from Hoechst+ cells based on high Hoechst intensity and extracellular DNA area (Sytox Orange staining beyond cell margins) across treatment groups. (**C**) Frequency of NET DNA fragments (Type IV) gated based on high extracellular DNA with lower Hoechst intensity. (**D**) Circularity score (degree of the Hoechst mask deviation from a circle, with lower values having more deviation) across treatment groups. (**E**) Additional representative images of Type III NETs from PMA and PMA + THP treatment groups showing variable levels of MPO staining across groups. Experiments were performed in four independent experiments with data combined, *n* = 4 donors. Box and whisker plots extend from 25th to 75th percentiles and show all points (A-D). Data were analyzed by one-way ANOVA with Holm-Sidak’s multiple comparisons test (A-D). * *P* < 0.05. Supplemental data to **Figure 7**.

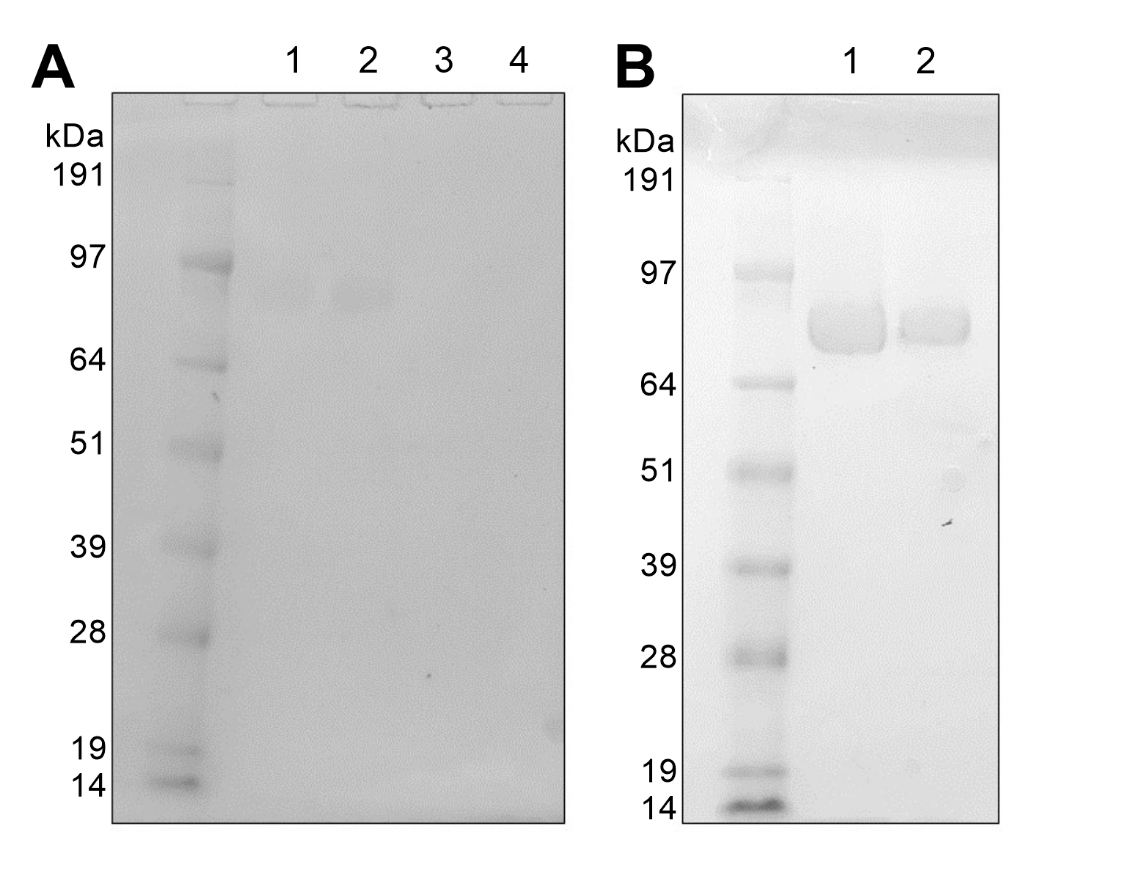

**Supplemental Figure 6. Representative protein gels of THP isolated from mouse and human urine.** (**A**) Polyacrylamide protein gel of THP isolated from pooled mouse urine as described in Methods. Lane 1: WT mock-infected, Lane 2: WT UPEC-infected, Lane 3: THP KO mock-infected, Lane 4: THP KO UPEC-infected. THP was visualized as a single band at ~ 85 kDa in lanes 1 and 2. (B) Polyacrylamide protein gel of THP isolated from pooled human urine as described in Methods. Lane 1: pooled batch #1, Lane 2: pooled batch #2. THP was visualized as a single band at ~ 85 kDa in lanes 1 and 2.

| **Supplemental Table 1. Comparative N-glycan MALDI-tof analyses of THP during UPEC infection.** | | | |
| --- | --- | --- | --- |
| m/z peak | WT Mock | WT UPEC | % change |
| 2244.459 | 1.93 | 1.31 | -47.1 |
| 2431.552 | 4.24 | 3.59 | -18.1 |
| 2605.648 | 8.40 | 9.00 | 6.6 |
| 2792.735 | 2.47 | 2.75 | 10.2 |
| 2966.841 | 14.28 | 17.17 | 16.9 |
| 3241.968 | 4.00 | 4.46 | 10.1 |
| 3416.062 | 8.41 | 13.05 | 35.6 |
| 3777.244 | 17.42 | 17.32 | -0.5 |
| 4226.453 | 14.22 | 16.24 | 12.5 |
| 4587.622 | 24.64 | 15.11 | -63.1 |

N-glycan MALDI-tof peaks of THP purified from urine of WT mice that were either mock-infected or UPEC-infected. Data represent one MALDI-tof analysis of pooled purified THP harvested from two independent experiments. Supplemental data to **Figure 3**.

**Supplemental table 2. Differentially abundant proteins between THP-treated and mock-treated human neutrophils in unstimulated and PMA-stimulated conditions.** Supplemental data to **Figure 5**.

| **Gene ID** | **Gene Symbol** | **Gene Description** | **Log_2_ Fold-Change Unstimulated^a^**  **(*p*-value)^c^** | **Log_2_ Fold-Change Stimulated^b^**  **(*p*-value)** |
| --- | --- | --- | --- | --- |
| 7369 | UMOD | uromodulin | 2.894  **(2.14E-12)** | 2.497  **(3.77E-11)** |
| 347 | APOD | apolipoprotein D | 2.112  **(2.14E-12)** | 1.200  **(6.47E-09)** |
| 55902 | ACSS2 | acyl-CoA synthetase short-chain family member 2 | 0.684  **(7.73E-08)** | 0.466  **(1.27E-05)** |
| 23506 | BICRAL | GLTSCR1-like | 0.669  **(0.039)** | 0.032  (0.962) |
| 3959 | LGALS3BP | lectin, galactoside-binding, soluble, 3 binding protein | 0.624  **(2.70E-09)** | 0.533  **(2.62E-08)** |
| 3827 | KNG1 | kininogen 1 | 0.612  **(7.79E-07)** | 0.485  **(1.58E-05)** |
| 2731 | GLDC | glycine dehydrogenase (decarboxylating) | 0.539  **(0.017)** | 0.445  (0.052) |
| 259 | AMBP | alpha-1-microglobulin/bikunin precursor | 0.488  **(3.02E-05)** | 0.647  **(1.47E-06)** |
| 220963 | SLC16A9 | solute carrier family 16, member 9 | 0.485  **(4.93E-04)** | 0.364  **(0.005)** |
| 3561 | IL2RG | interleukin 2 receptor, gamma | 0.456  **(3.02E-05)** | 0.325  **(0.001)** |
| 10621 | POLR3F | polymerase (RNA) III (DNA directed) polypeptide F, 39 kDa | 0.422  **(0.030)** | 0.184  (0.445) |
| 952 | CD38 | CD38 molecule | 0.382  **(0.017)** | 0.142  (0.487) |
| 55647 | RAB20 | RAB20, member RAS oncogene family | 0.554  **(0.046)** | 0.088  (0.848) |
| 440275 | EIF2AK4 | eukaryotic translation initiation factor 2 alpha kinase 4 | 0.352  **(1.66E-06)** | 0.201  (0.001) |
| 107987285 | LOC107987285 | uncharacterized protein | 0.344  **(0.042)** | 0.030  (0.928) |
| 8225 | GTPBP6 | GTP binding protein 6 (putative) | 0.221  (0.380) | 0.433  **(0.030)** |
| 1521 | CTSW | cathepsin W | 0.216  (0.605) | -0.704  **(0.011)** |
| 3576 | CXCL8 | chemokine (C-X-C motif) ligand 8 | 0.204  (0.409) | 0.406  **(0.038)** |
| 54810 | GIPC2 | GIPC PDZ domain containing family, member 2 | 0.167  (0.523) | 0.410  **(0.030)** |
| 9698 | PUM1 | pumilio RNA-binding family member 1 | 0.160  (0.515) | 0.392  **(0.028)** |
| 722 | C4BPA | complement component 4 binding protein, alpha | 0.109  (0.807) | -0.529  **(0.019)** |
| 91272 | BOD1 | biorientation of chromosomes in cell division 1 | 0.104  (0.666) | 0.354  **(0.018)** |
| 678 | ZFP36L2 | ZFP36 ring finger protein-like 2 | 0.103  (0.510) | 0.357  **(0.002)** |
| 8031 | NCOA4 | nuclear receptor coactivator 4 | 0.055  (0.805) | 0.328  **(0.003)** |
| 966 | CD59 | CD59 molecule, complement regulatory protein | 0.050  (0.760) | 0.369  **(1.42E-04)** |
| 1774 | DNASE1L1 | deoxyribonuclease I-like 1 | 0.045  (0.768) | 0.332  **(2.02E-04)** |
| 5150 | PDE7A | phosphodiesterase 7A | 0.041  (0.906) | 0.425  **(0.003)** |
| 79080 | CCDC86 | coiled-coil domain containing 86 | 0.024  (0.974) | 0.545  **(0.030)** |
| 1871 | E2F3 | E2F transcription factor 3 | -0.018  (0.958) | 0.400  **(0.002)** |
| 197258 | FCSK | fucokinase | -0.069  (0.713) | 0.385  **(0.001)** |
| 129446 | XIRP2 | xin actin binding repeat containing 2 | -0.085  (0.768) | -0.528  **(0.001)** |
| 7915 | ALDH5A1 | aldehyde dehydrogenase 5 family, member A1 | -0.348  **(0.031)** | -0.094  (0.696) |
| 10422 | UBAC1 | UBA domain containing 1 | -0.378  **(0.017)** | 0.048  (0.861) |
| 10193 | RNF41 | ring finger protein 41, E3 ubiquitin protein ligase | -0.482  **(0.038)** | -0.104  (0.783) |

^a^Fold-change between THP-treated and mock-treated samples in unstimulated conditions.

^b^Fold-change between THP-treated and mock-treated samples in PMA-stimulated conditions.

^c^Differential proteins were identified via Log_2_ fold change >1.25 and moderated t-test followed by multiple-hypothesis testing correction using the Benjamini–Hochberg procedure with a false discovery rate adjusted *P* <0.05. Proteins meeting these criteria are indicated by p-values in bold text.
